# The WalRK two-component system is essential for proper cell envelope biogenesis in *Clostridioides difficile*

**DOI:** 10.1101/2022.04.01.486798

**Authors:** Ute Müh, Craig D. Ellermeier, David S. Weiss

## Abstract

The WalR-WalK two-component regulatory system (TCS) is found in all Firmicutes, where it regulates the expression of multiple genes required for remodeling the cell envelope during growth and division. Unlike most TCSs, WalRK is essential for viability, so it has attracted interest as a potential antibiotic target. Here we used overexpression of WalR and CRISPR interference to investigate the Wal system of *Clostridioides difficile*, a major cause of hospital-associated diarrhea in high-income countries. We confirmed the *wal* operon is essential and identified morphological defects and cell lysis as the major terminal phenotypes of altered *wal* expression. We also used RNA-seq to identify over 150 genes whose expression changes in response to WalR levels. This gene set is enriched in cell envelope genes and includes several predicted PG hydrolases and proteins that could regulate PG hydrolase activity. A distinct feature of the *C. difficile* cell envelope is the presence of an S-layer and we found WalR affects expression of several genes which encode S-layer proteins. An unexpected finding was that some Wal-associated phenotypic defects were inverted in comparison to what has been reported in other Firmicutes. For example, down-regulation of Wal signaling caused *C. difficile* cells to become longer rather than shorter, as in *Bacillus subtilis*. Likewise, down-regulation of Wal rendered *C. difficile* more sensitive to vancomycin, whereas reduced Wal activity is linked to increased vancomycin resistance in *Staphylococcus aureus*.

**Importance:** The WalRK two-component system (TCS) is essential for coordinating synthesis and turnover of peptidoglycan in Firmicutes. Here we investigate the WalRK TCS in *Clostridioides difficile*, an important bacterial pathogen with an atypical cell envelope. We confirmed that WalRK is essential and regulates cell envelope biogenesis, although several of the phenotypic changes we observed were opposite to what has been reported in other Firmicutes. We also identified over 150 genes whose expression is controlled either directly or indirectly by WalR. Overall, our findings provide a foundation for future investigations of an important regulatory system and potential antibiotic target in *C. difficile*.

## Introduction

To grow and divide, bacteria must thoroughly remodel their peptidoglycan (PG) sacculus, a large, covalently closed macromolecule that surrounds the cell and provides protection against rupture due to turgor pressure (1). How bacteria remodel the sacculus without making errors that lead to inadvertent lysis is a question that has fascinated microbiologists for decades. The question is also of practical significance because small molecules that undermine normal peptidoglycan biogenesis are among our most effective antibiotics.

An important advance in our understanding of cell envelope biogenesis was made over twenty years ago when Fabret and Hoch identified a *Bacillus subtilis* two-component system (TCS) that is essential for viability (2). This TCS is now called the Wal system and is known to coordinate expression of multiple cell envelope genes in the Firmicutes. WalRK is essential in all species studied so far, including pathogens such as *Staphylococcus aureus* (3–5) and *Streptococcus pneumoniae* (6, 7). Because of its essentiality, the Wal TCS has attracted interest as a potential antibiotic target.

The Wal TCS comprises the bifunctional kinase/phosphatase WalK together with its cognate response regulator WalR. In addition, *wal* operons typically include genes for membrane proteins known or presumed to modulate WalK activity, but the accessory proteins differ between organisms (8–10). Recent evidence from *B. subtilis* indicates WalK activity is regulated by PG fragments generated by PG hydrolases that open spaces to make room for elongation. These PG fragments bind to WalK’s extracellular Cache domain to modulate the phosphorylation state of WalR (11). Phosphorylated WalR in turn binds to various promoters to activate or repress gene expression.

The number of genes in the WalR regulon ranges from about a dozen to over 100 (5, 12–16), depending on the species, and only some of these genes are directly activated or repressed by WalR. Although WalR regulons are diverse, they invariably include multiple proteins that contribute to proper biogenesis of the cell envelope, especially PG hydrolases. Consistent with these themes, the phenotypic defects elicited by artificial up- or down-regulation of WalRK signaling include abnormal cell shape, larger cells, smaller cells, ghost cells (lysis), thicker PG, abnormal division septa and altered sensitivity to antibiotics that target PG biogenesis (8–10). In *S. pneumoniae walRK* is essential because it activates expression of an essential PG hydrolase, *pcsB* (6, 7). However, the essentiality of *walRK* in *B. subtilis* and *S. aureus* is polygenic in nature, resulting from abnormal expression of multiple genes that are not by themselves essential. In all three organisms, it is possible to bypass the *walRK* requirement by constitutively expressing one or more Wal regulon PG hydrolases, sometimes in combination with deleting genes for hydrolase inhibitors (6, 17, 18).

Here we studied the WalRK TCS in *Clostridioides difficile*, an anaerobic, spore-forming Firmicute responsible for close to a quarter million hospitalizations and over 12,000 deaths per year in the United States (19). Unlike the envelope of most Firmicutes in which *walRK* has been studied previously, the *C. difficile* envelope has a proteinaceous S-layer whose assembly presumably must be coordinated with PG synthesis (20–22). In addition, the PG itself is somewhat unusual (23). About 90% of the *N*-acetylglucosamine is deacetylated (which has implications for PG hydrolase activity) and about 70% of the peptide crosslinks are 3-3 rather than 4-3 (which has implications for the structure of PG fragments thought to bind to the Cache domain of WalK (11)). The *C. difficile* Wal system includes a unique lipoprotein gene and a non-canonical WalK that lacks an intracellular PER-ARNT-SIM (PAS) signaling domain (Fig. 1) (10). These features distinguish *C. difficile’s* WalRK TCS from those of previously characterized Firmicutes, although a distantly related *wal*-like system in *Mycobacterium tuberculosis* (MtrAB) includes a lipoprotein and a WalK-like histidine kinase without an intracellular PAS domain (10, 24, 25). Finally, our recent development of tools for xylose-inducible gene expression and CRISPR interference (26) helped to overcome some of the challenges inherent in phenotypic analysis of essential genes like *walRK* in *C. difficile*.

**Fig. 1.**
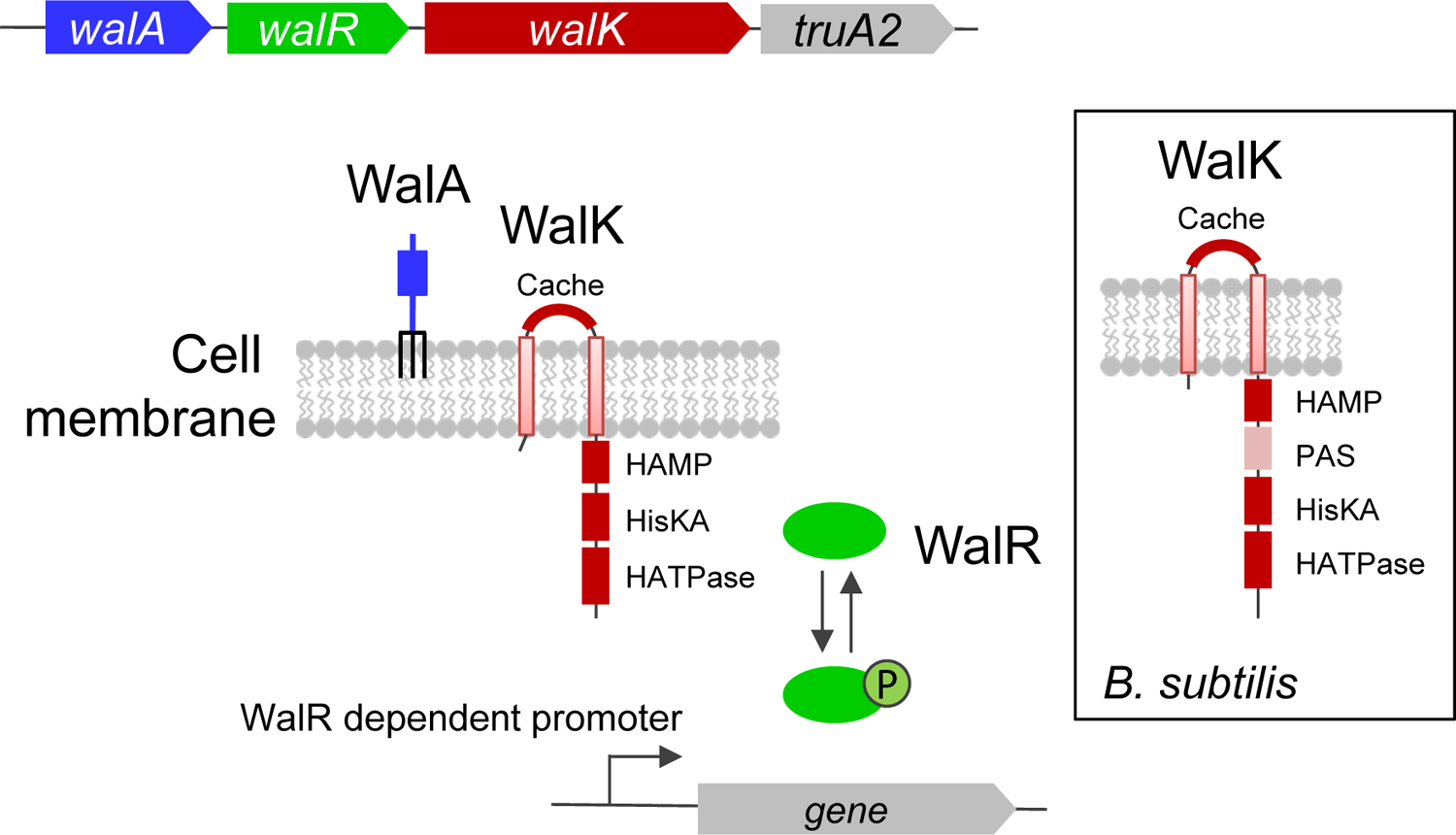
The *wal* operon of *C. difficile*. Based on studies in other bacteria, WalK (red) is a bifunctional signal transducing enzyme that can phosphorylate or dephosphorylate WalR (green). WalK is predicted to have an extracellular Cache domain, as well as intracellular HAMP, HisKA (phospho-acceptor), and HATPase domains. Unlike the *B. subtilis* ortholog (inset), it does not have an intracellular PAS domain. WalR∼P binds DNA to activate or repress expression of Wal-regulated genes. The lipoprotein (WalA, blue), which is unique to *C. difficile*, may modulate WalK activity and is predicted to have a beta-propeller lyase domain. The *truA2* gene codes for a pseudouridylate synthase and is not thought to play a role in Wal signaling. Operon locus: *cdr20291_1676-1679* in R20291 and *cd630_17810-17840* in 630Δerm.

## Results

### The *wal* operon is required for cell viability, proper rod morphology and intrinsic resistance to some antibiotics

According to Tn-seq, the WalRK TCS is essential in *C. difficile* (27). To confirm and extend this finding, we leveraged the power of CRISPR interference (CRISPRi) for functional analysis of essential genes/operons (28). Our CRISPRi system comprises a xylose-inducible nuclease-defective Cas9, P*_xyl_*::*dCas9*, that is targeted to a gene of interest by a single guide RNA expressed constitutively from the glutamate dehydrogenase promoter, P*_gdh_*::*sgRNA* (26). Because CRISPRi is polar, we decided to target the first gene in the operon, which encodes a predicted lipoprotein unique to the *C. difficile wal* operon (Fig. 1). We have named this gene *walA.* For reproducibility, we designed two guides against *walA*, sgRNA-*walA1* and sgRNA-*walA2*. Subsequent RNA-seq experiments described below confirmed that targeting dCas9 to *walA* suppressed transcription of the entire operon. As a negative control we used a sgRNA (sgRNA-neg) that does not have a target anywhere on the *C. difficile* chromosome. In some experiments the CRISPRi machinery was produced from a plasmid that confers resistance to thiamphenicol (Thi), while in others it was integrated into the chromosome at the *pyrE* locus in a way that does not leave behind an antibiotic resistance marker (29).

Chromosomal- or plasmid-based CRISPRi knockdown of the *wal* operon in either 630Δerm or R20291 reduced viability ≥10^4^-fold when cell were plated on TY-agar containing 1% xylose (Fig. 2A and Fig. S1A). Additional phenotypes were characterized in TY broth using 630Δerm derivatives with the CRISPRi machinery integrated at *pyrE*. In the absence of xylose, sgRNA-*walA1* and sgRNA-*walA2* retarded growth slightly compared to the sgRNA-neg control (Fig. 2B, S1B). This finding indicates there is leaky expression of P*_xyl_*::*dCas9,* which was later confirmed by RNA-seq (see below). Under these conditions, cells were on average about 10% longer than wild-type and the fraction of cells with curved or irregular contours increased from ∼1% in the sgRNA-neg control to ∼20% for sgRNA-*walA1* or ∼10% for sgRNA-*walA2* (Fig. 2C-E, S1E,F). Further knockdown of the *wal* operon by addition of xylose exacerbated the growth defect in a dose-dependent manner (Fig. 2B, S1D). In addition, morphological defects became more pronounced, with cells now averaging ∼30% longer than controls and up to ∼40% exhibiting curved or irregular contours (Fig. 2D,E and S1E,F).

**Fig. 2.**
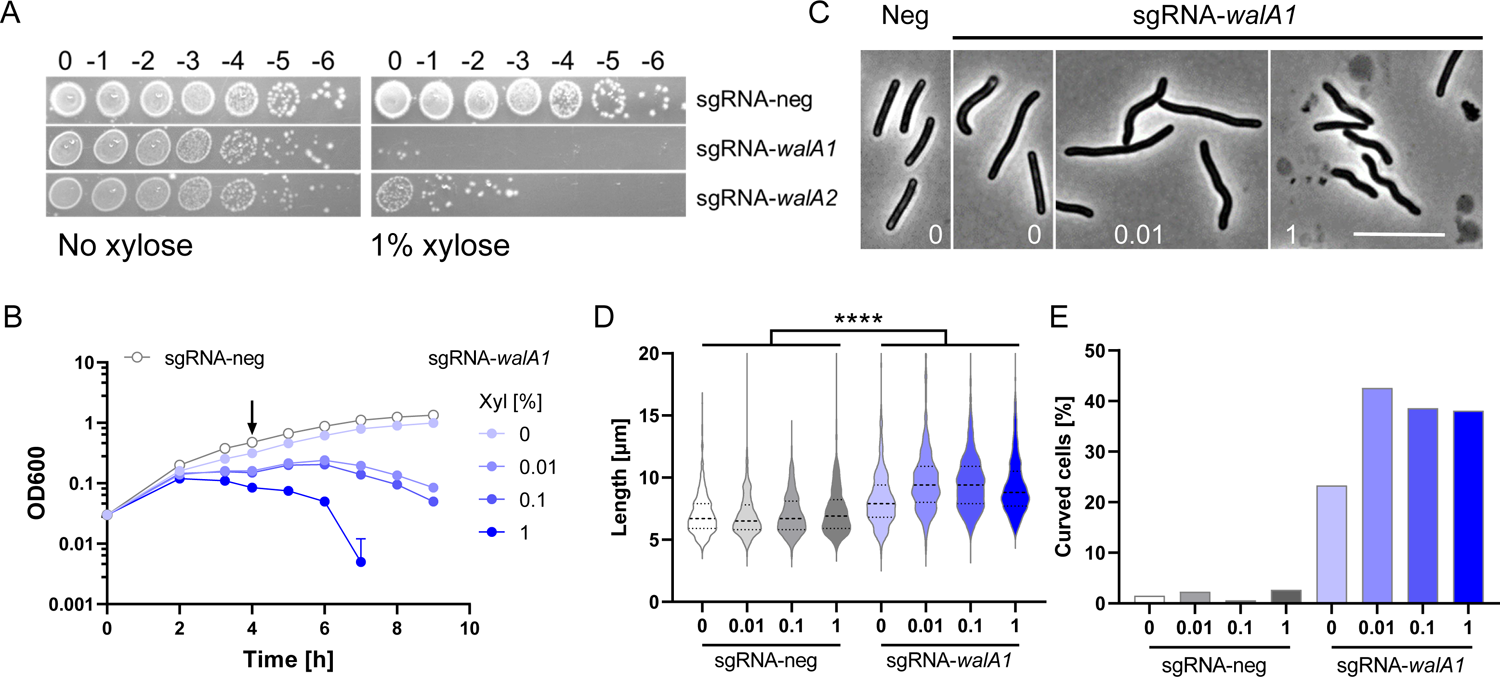
The *wal* operon is required for viability and normal rod morphology of *C. difficile* 630Δerm. (A) Viability assay. Serial dilutions of overnight cultures were spotted onto TY plates with or without 1% xylose to induce the expression of *dCas9*. Plates were photographed after incubation overnight. Strains carry chromosomal copies of the CRISPRi components and the following guides: sgRNA-neg (UM554), sgRNA-*walA1* (UM555), and sgRNA-*walA2* (UM556). (B) Growth curves in liquid TY containing 0, 0.01, 0.1 or 1% xylose as indicated. For clarity, the sgRNA-neg control strain is graphed only at 0 xylose. Samples were taken at 4 hours (arrow) for phase contrast microscopy (C), cell length measurements (D) and determination of curvature (E). Numbers at the bottom of micrographs in (C) refer to % xylose. Note cell debris indicative of lysis at 1% xylose. Bar = 10 µm. Cell length and percent curved cells were based on about 500 cells per condition. The sgRNA-*walA1* strain is longer than the sgRNA-neg control in all pairwise comparisons as determined by a T-test, p<0.0001.

Partial knockdown of the *wal* operon in the absence of xylose allowed us to screen for altered sensitivity to antibiotics that target cell envelope biogenesis. We found that the minimal inhibitory concentration (MIC) for ampicillin and daptomycin decreased about 2-fold, while the MIC for vancomycin decreased about 4-fold (Table 1). But sensitivity to the other two cell wall-active antibiotics, imipenem and bacitracin, or the gyrase inhibitor novobiocin was not affected (Table 1).

**Table 1.**
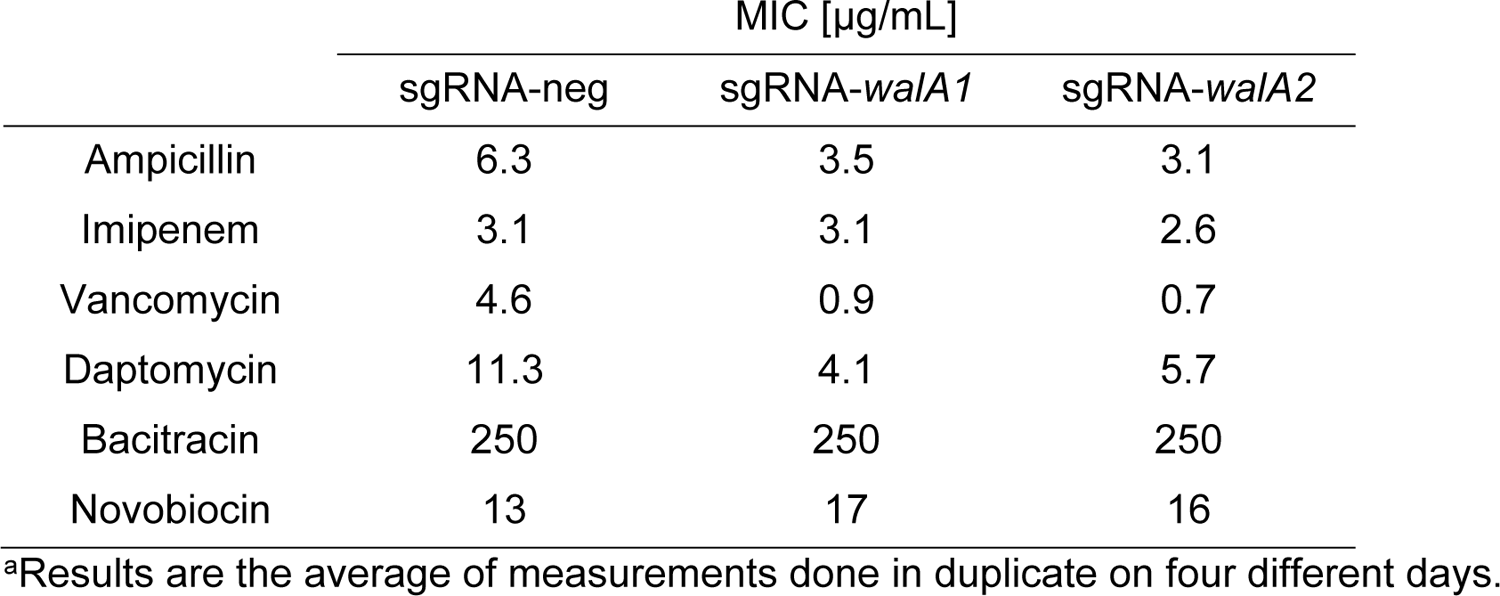
Effect of CRISPRi-silencing *wal* on sensitivity to antibiotics^a^

### Overexpression of WalR impairs growth, alters morphology and slows autolysis

We also investigated Wal function in *C. difficile* by overproducing the response regulator WalR to drive cells into an exacerbated Wal-ON state (5, 30). Plating *C. difficile* 630Δerm harboring a P*_xyl_*::*walR* expression plasmid onto TY-Thi containing 1% xylose resulted in a 10^6^-fold loss of viability (Fig. 3A). This strain failed to grow when subcultured directly into TY-Thi broth containing 1% xylose, but waiting until OD_600_ ∼0.2 before adding 3% xylose resulted in only a subtle growth defect (Fig. 3B). Three hours post-induction, the cells had become strikingly phase bright and resistant to autolysis when suspended in buffer containing 0.01% Triton X-100 (Fig. 3C,D). Triton X-100 is thought to reduce the inhibitory effect of lipoteichoic acids on PG hydrolases (31, 32). Similar results were obtained in the R20291 strain background (Fig. S3). It is unclear what causes cells to turn phase bright upon overexpression of *walR*. Differential staining with Syto 9 and propidium iodide (LIVE/DEAD staining) indicates that >99% of the cells are alive after overproduction of WalR (Fig. 3B, red arrow).

**Fig. 3.**
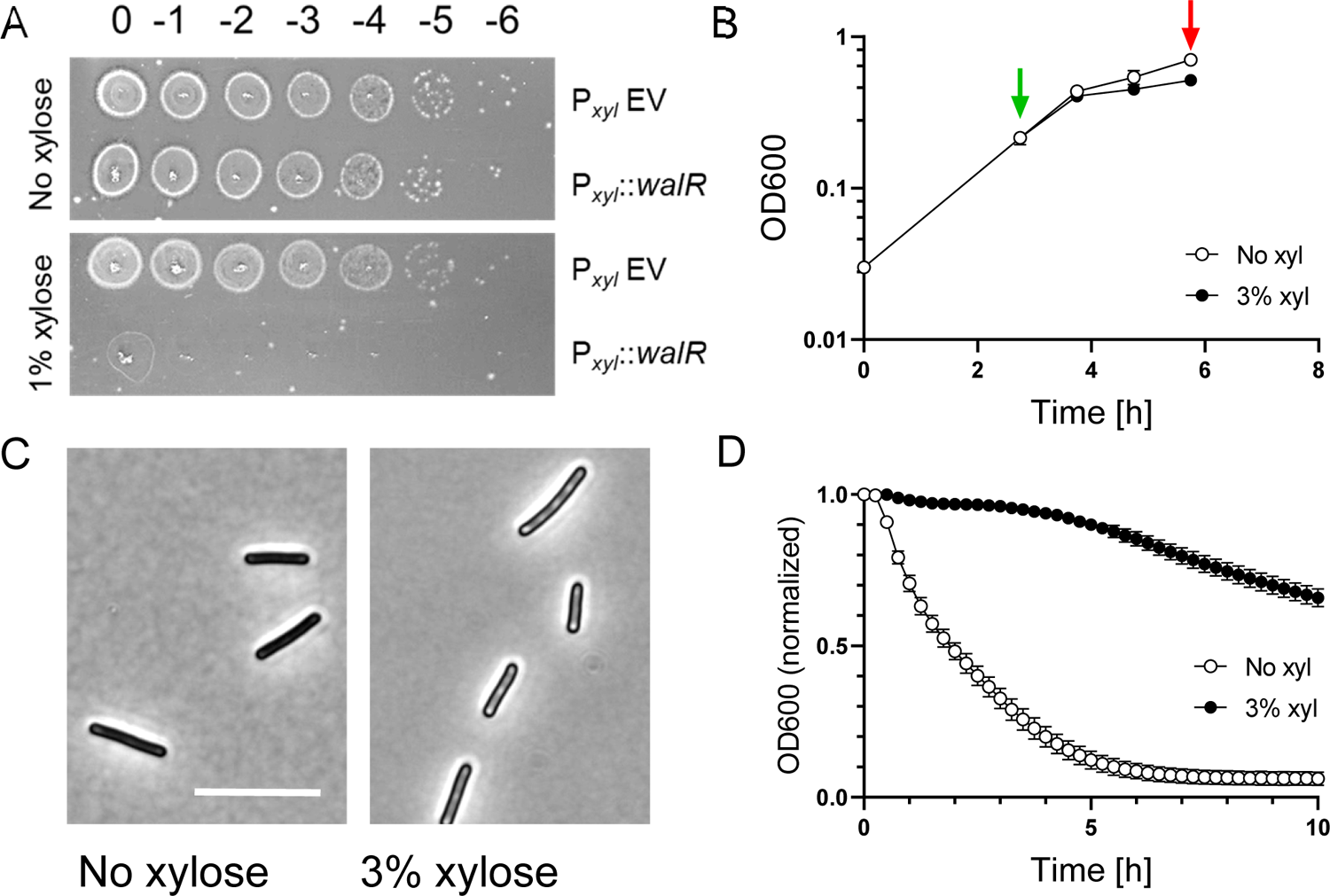
Overexpression of *walR* from a multi-copy plasmid impairs growth, alters morphology and slows autolysis in 630Δerm. A. Viability assay. Overnight cultures of 630Δerm harboring pCE691 (P*_xyl_*::*walR*) or the empty vector control plasmid pBZ101 (P*_xyl_* EV) were serially diluted and spotted onto TY-Thi plates with or without 1% xylose. Plates were photographed after incubation overnight. (B) Growth curve of 630Δerm/pCE691 (P*_xyl_*::*walR*). Duplicate cultures were grown to an OD_600_ of 0.2, at which time one was induced with 3% xylose (green arrow). Both cultures were harvested after 3h (red arrow) for analysis by phase contrast microscopy (C) and a lysis assay (D). Bar in (C) = 10 µm. For the lysis assay, cells were suspended in buffer containing 0.01% Triton X-100 and OD_600_ was monitored over time. Error bars depict SD of 3 technical replicates. Data shown are representative of at least 3 independent experiments.

We repeated these experiments after integrating P*_xyl_::walR* into the chromosome of 630Δerm. As expected, the effects of xylose induction were similar but less pronounced when P*_xyl_::walR* was in single copy (Fig. 4A). The addition of xylose at subculture was now tolerated and reduced growth in a dose-dependent manner. At 3% xylose, cells became slightly phase-bright. There were some bent or hooked cells, a defect not observed with the P*_xyl_::walR* plasmid, perhaps because it could only be induced for shorter times. (Fig. 4C). After 6h of induction, cells were harvested and evaluated in the lysis assay. We found that overexpression of *walR* slowed lysis in a dose-dependent manner (Fig. 4B).

**Fig. 4.**
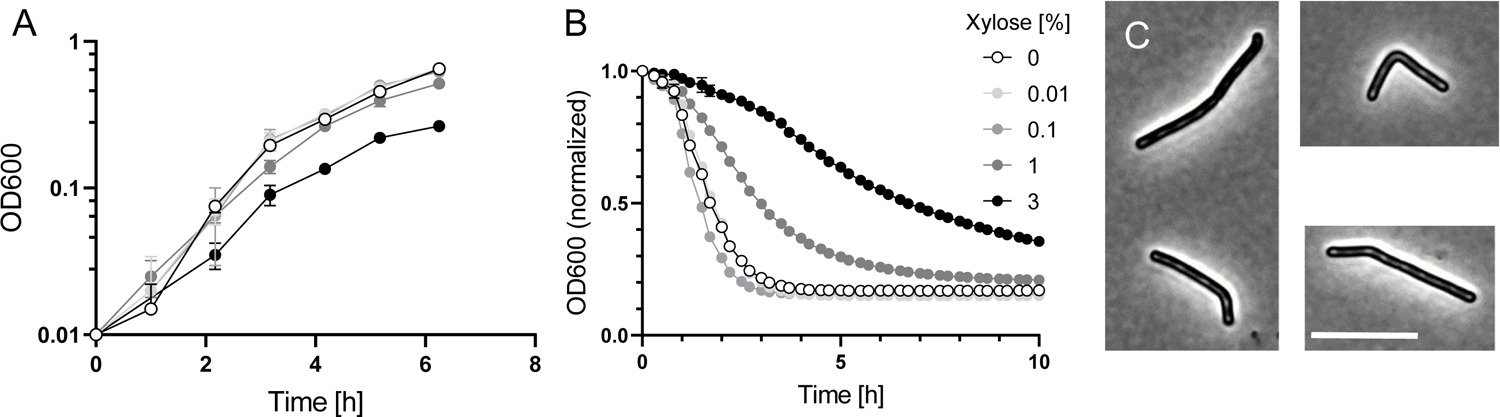
Prolonged overexpression of *walR* from a single-copy chromosomal P*_xyl_*::*walR* allele affects cell shape and slows lysis. (A) Growth curves. An overnight culture of strain UM626 (P*_xyl_::walR* integrated at *pyrE* in 630Δerm) was subcultured 1:50 into TY with varied xylose concentrations as indicated and grown for 6 hours, then analyzed in a lysis assay (B) or by phase contrast microscopy (C). For the lysis assay, cells were suspended in buffer containing 0.01% Triton X-100 and OD_600_ was monitored over time. Data are graphed as the mean and SD of 3 technical replicates, but most error bars are smaller than the symbols. Cells from cultures grown in the presence of 0-1% xylose appeared normal and are not shown, but in cultures induced with 3% xylose about 10% of the cells had irregular or bent morphologies, examples of which are shown in (C). Bar = 10 μm. Data shown are representative of at least 3 independent experiments.

### Expression profiling under Wal-ON conditions

As a step towards understanding the basis of the various Wal-related phenotypes, we sought to identify the genes in the *wal* regulon. (Note: we use the term regulon for convenience to refer to genes whose expression responds to manipulation of the *wal* operon regardless of whether these effects are direct or indirect.) We began by performing RNA-seq on two strains that overexpress *walR* from a single copy P*_xyl_*::*walR* construct integrated at *pyrE*. One of these strains overproduced wild-type WalR, the other overproduced a WalR^D54E^ mutant protein expected to mimic phosphorylated WalR and thus be constitutively active (13, 33) (Confusingly, D54 is annotated as D66 in the R20291 genome on the BioCyc website). The two strains behaved similarly with respect to growth and performance in the lysis assay (Fig. S4A,B). The control strain for these experiments had the gene for a red fluorescent protein (*rfp*) integrated at *pyrE* (26). Strains were grown overnight in TY, then subcultured into fresh TY at a starting OD_600_ ∼0.05. When the OD_600_ reached ∼0.2, cultures were induced with 3% xylose and allowed to grow for three hours (∼ 2 mass doublings), at which time cells were harvested and processed for RNA-seq. At this time point induction of *walR* had only a small effect on growth rate (Fig. S4A) and >99% were viable as judged by LIVE/DEAD staining.

Not including *walR*, which was induced directly from P*_xyl_*, the RNA-seq analysis identified 77 genes whose transcript abundance changed ≥4-fold (p<0.05) upon overexpression of either *walR* or *walR*^D54E^ (Table 2). Of these genes, 44 were induced and 33 were repressed. Induction ratios were very similar in the *walR* and *walR^D54E^* strains with only two exceptions, *cd630_03901* and *cd630_10631*, both of which encode hypothetical proteins. The failure of the D54E substitution to increase gene expression could mean it is not activating in *C. difficile* WalR, as reported previously for some other response regulators (34). Alternatively, overexpression of *walR* is itself activating and might obscure the effects of the substitution. The 77 genes are predicted by BioCyc (35) to be organized into 65 operons that contain 18 additional genes which missed our 4-fold expression change cut-off. Visual inspection of the expression data revealed these genes almost always trended in the right direction, confirming the predicted structure of most of the operons. We therefore included these 18 genes in what we will refer to as the Wal-ON regulon, bringing it to a total of 95 genes in 65 operons (Table 2).

**Table 2.**
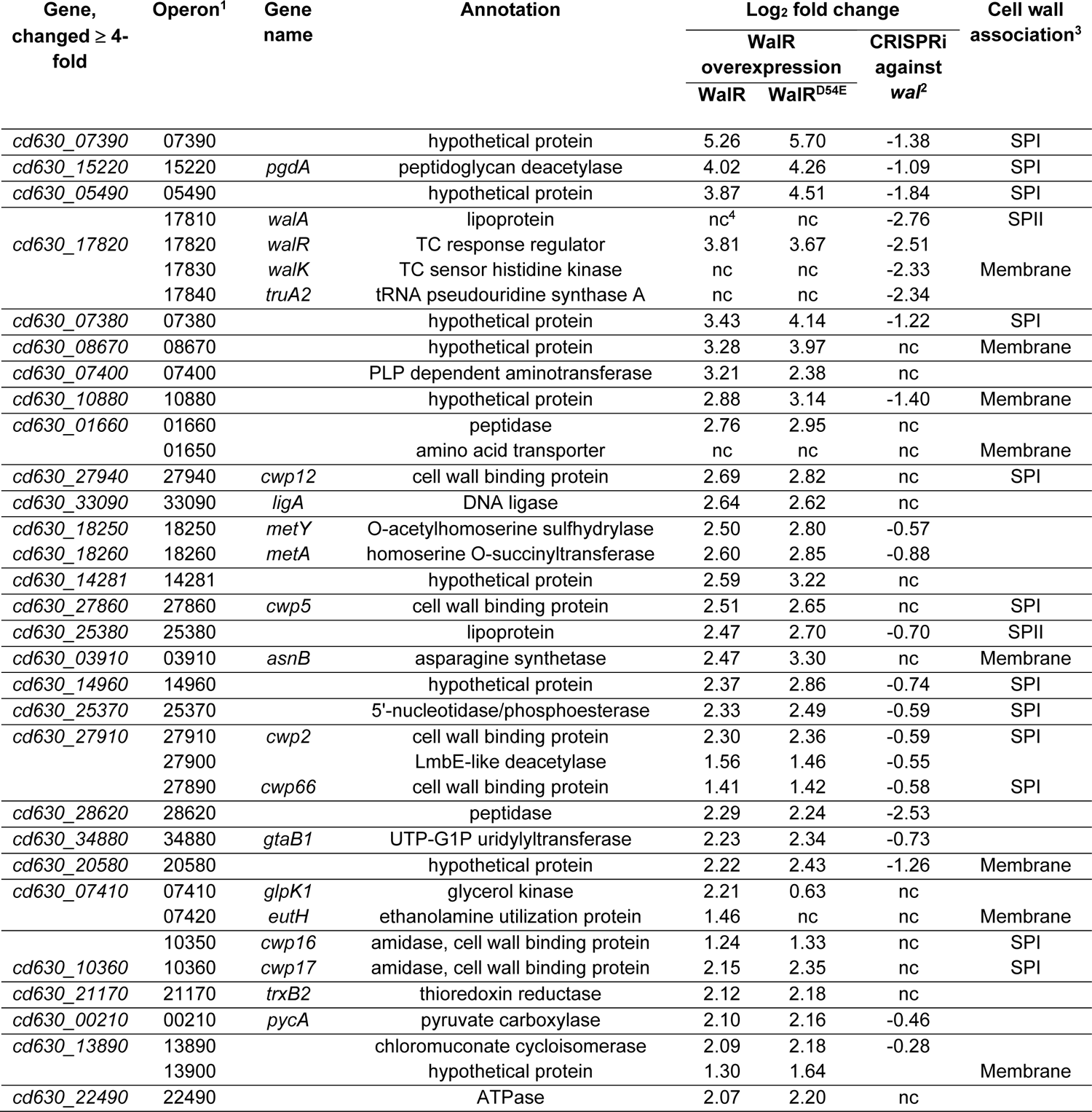
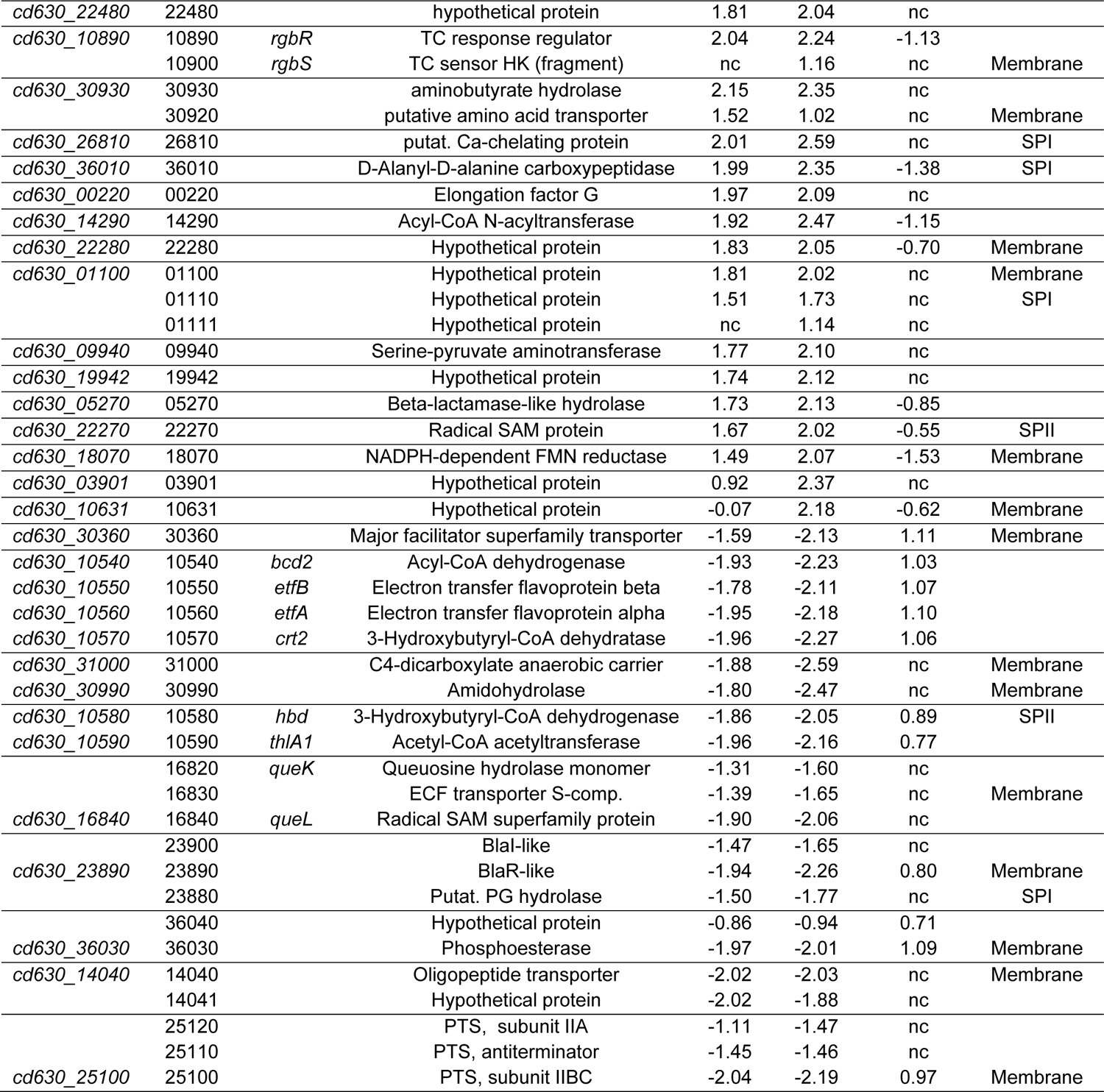
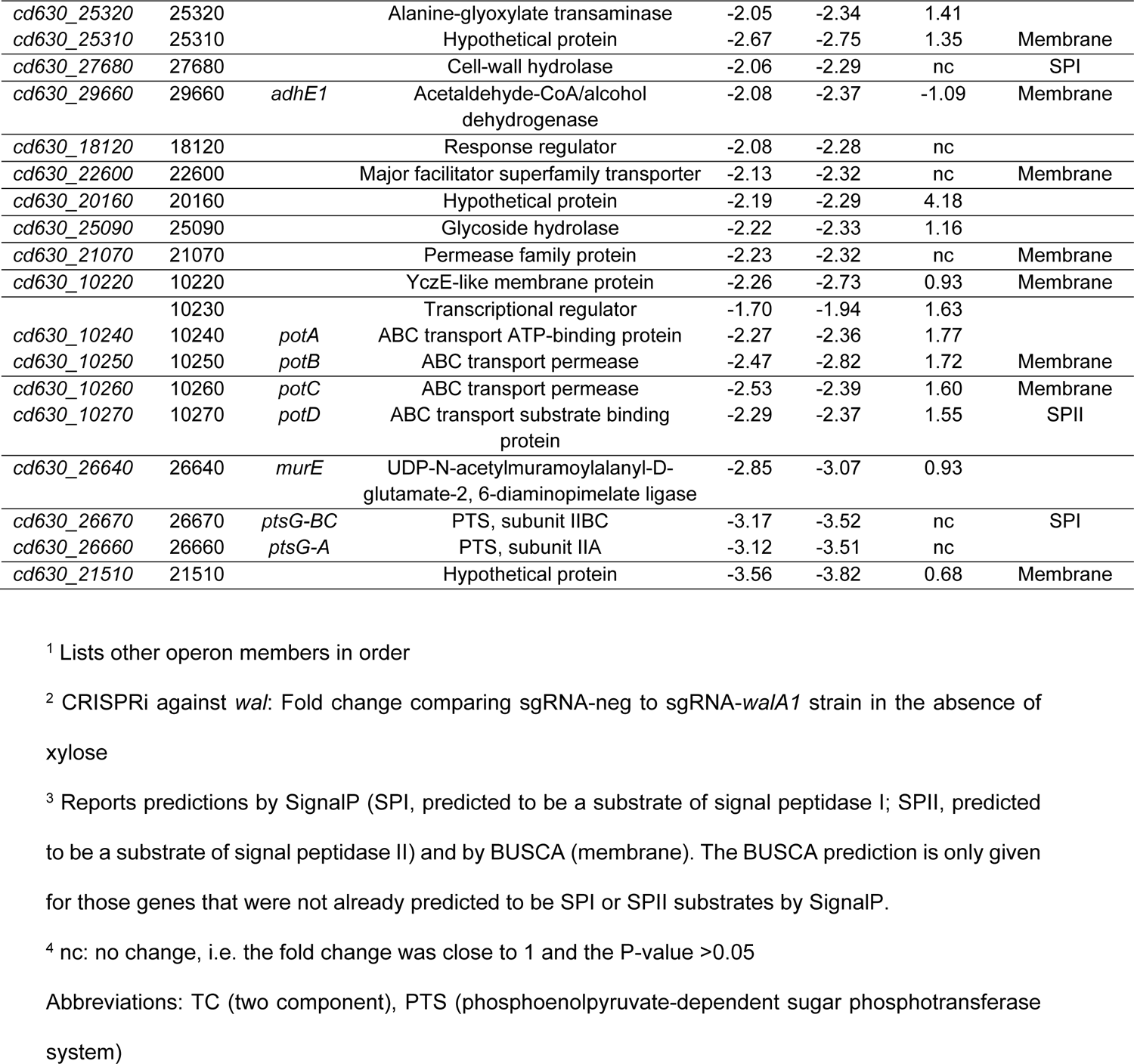
Genes affected by Wal-ON conditions.

Several of the genes and trends in Table 2 are worth highlighting. Xylose induction resulted in a 14-fold increase in *walR* mRNA (13-fold for *walR*^D54E^), while the levels of *walK* or other members of the native operon were not affected. This means that induction of P*_xyl_*::*walR* integrated at *pyrE* does not lead to induction of the native *wal* operon, consistent with reports that the *wal* operon is not auto-regulated in other bacteria (8, 16). Another striking feature of Table 2 is the abundance of genes encoding for cell envelope proteins. Of the 95 genes in Table 2, 22 (23%) are predicted to encode exported proteins and 29 (31%) are predicted to encode cytoplasmic membrane proteins. For comparison, ∼7% of the entire proteome is predicted to be exported and 23% predicted to be membrane proteins (see Methods). Table 2 also includes 21 hypotheticals (22% of the total), many of which are likely to be involved in cell envelope processes based on the presence of predicted transmembrane domains or export signals.

Peptidoglycan-associated genes, especially PG hydrolases, are prominent members of the Wal regulon in all Firmicutes examined to date (8, 10), and *C. difficile* is no exception. Table 2 lists ten peptidoglycan-associated genes, including the second most highly induced gene, *pgdA*, which was upregulated 16-fold. PgdA deacetylates N-acetylglucosamine in peptidoglycan, and is important for *C. difficile’s* high lysozyme resistance (36, 37). AsnB (∼8-fold induced) is annotated as an asparagine synthetase, but its primary function is to amidate diaminopimelic acid in PG stem peptides (38). In *B. subtilis*, amidation of m-DAP inhibits PG hydrolase activity (39). MurE (7-fold repressed) catalyzes one of the cytoplasmic steps in PG synthesis, addition of diaminopimelic acid to the nucleotide-linked PG precursor (40). Down regulation of *murE* suggests PG synthesis may be decreased in Wal-ON conditions. Cwp16 and Cwp17 are S-layer proteins with predicted amidase domains (41) and were upregulated about 4-fold. Cell wall amidases cleave the amide bond that links stem peptides to N-acetylglucosamine in PG glycan strands (42). CD630_36010 (4-fold induced) is a predicted D-alanyl-D-alanine carboxypeptidase that removes the terminal D-Ala moiety from peptidoglycan pentapeptide sidechains. This might limit 4-3 crosslinking of stem peptides, but could favor 3-3 crosslinking. Two proteins of unknown function were upregulated >10-fold, CD630_07390 and CD630_07380. We suspect both of these proteins are involved in PG metabolism because they are predicted to be exported and also induced by lysozyme (43). Notable among the repressed genes in Wal-ON cells is *cd630_27680*, which is down regulated about 4-fold and predicted to encode a cell wall hydrolase from the NlpC/P60 family, a domain common to endopeptidases (42). Interestingly, this gene is preceded by two putative WalR binding sites, and it has also been characterized as a sortase substrate (44). Another predicted PG hydrolase, CD630_23880, is mildly repressed (∼3-fold). Interestingly, this gene is cotranscribed with an uncharacterized BlaI-BlaR regulatory system, *cd630_23890* and *cd630_23900*. Some BlaIR regulatory systems respond to beta-lactam stress (45, 46).

One of the questions that motivated this study was whether the WalR system regulates genes for S-layer proteins in *C. difficile*. Indeed, the Wal-ON regulon includes six S-layer protein genes, all of which are upregulated by WalR: *cwp2*, *5*, *12*, *16*, *17* and *66*. Two of these genes were mentioned already, the cell wall amidases *cwp16* and *cwp17*. The remaining four are of unknown function, although *cwp2* and *cwp12* have domains implicated in adhesion or pathogenesis, and studies with *cwp66* have linked it to adhesion, autolysis and resistance to antibiotics (41, 47). WalRK has been suggested to regulate an S-layer gene in *Bacillus anthracis* (48).

### Expression profiling under Wal-OFF conditions

As a complementary approach to identify genes in the WalR regulon, we assessed the effect of CRISPRi knockdown of the *wal* operon on global gene expression. To this end, we performed RNA-seq on a 630Δerm derivative that has the CRISPRi machinery (P*_xyl_*::*dCas9*, P*_gdh_*::*sgRNA-walA1*) integrated into the chromosome at *pyrE*. The control strain was identical except that it expressed an innocuous sgRNA that does not target anywhere in the *C. difficile* genome (P*_gdh_*::*sgRNA-*neg). Two induction conditions were used. One of these, called “no xylose”, relied on leaky expression of *dCas9*. For this, overnight cultures were diluted into fresh TY to a starting OD_600_ = 0.03 and harvested when they reached OD_600_ ∼0.7 (Fig. S4C). For the other condition, called “low xylose”, cultures in exponential growth where induced with 0.1% xylose at an OD_600_ = 0.1 and incubated 2.5 hours (∼2 mass doublings) before harvest. This induction regimen reduced cell density only modestly (Fig. S4D).

There are 49 genes, organized in 27 operons that shift 4-fold or more under Wal-OFF conditions (Table 3). The 27 operons contain an additional 14 genes that did not meet our 4-fold cut-off, all but three of which nevertheless trended in the right direction. The three exceptions include a pseudogene (*cd630_05260*) that is not detected by our RNA-seq analysis tool and two membrane proteins (*cd630_05290* and *05280*) whose expression was unchanged. Thus, for the purposes of this analysis, CRISPRi knockdown of the *wal* operon changed the expression of 60 genes, of which 44 were induced and 16 were repressed. Curiously, for most of these genes the fold-change was larger in the absence of xylose, suggesting that many of the effects on gene expression might be indirect.

**Table 3.**
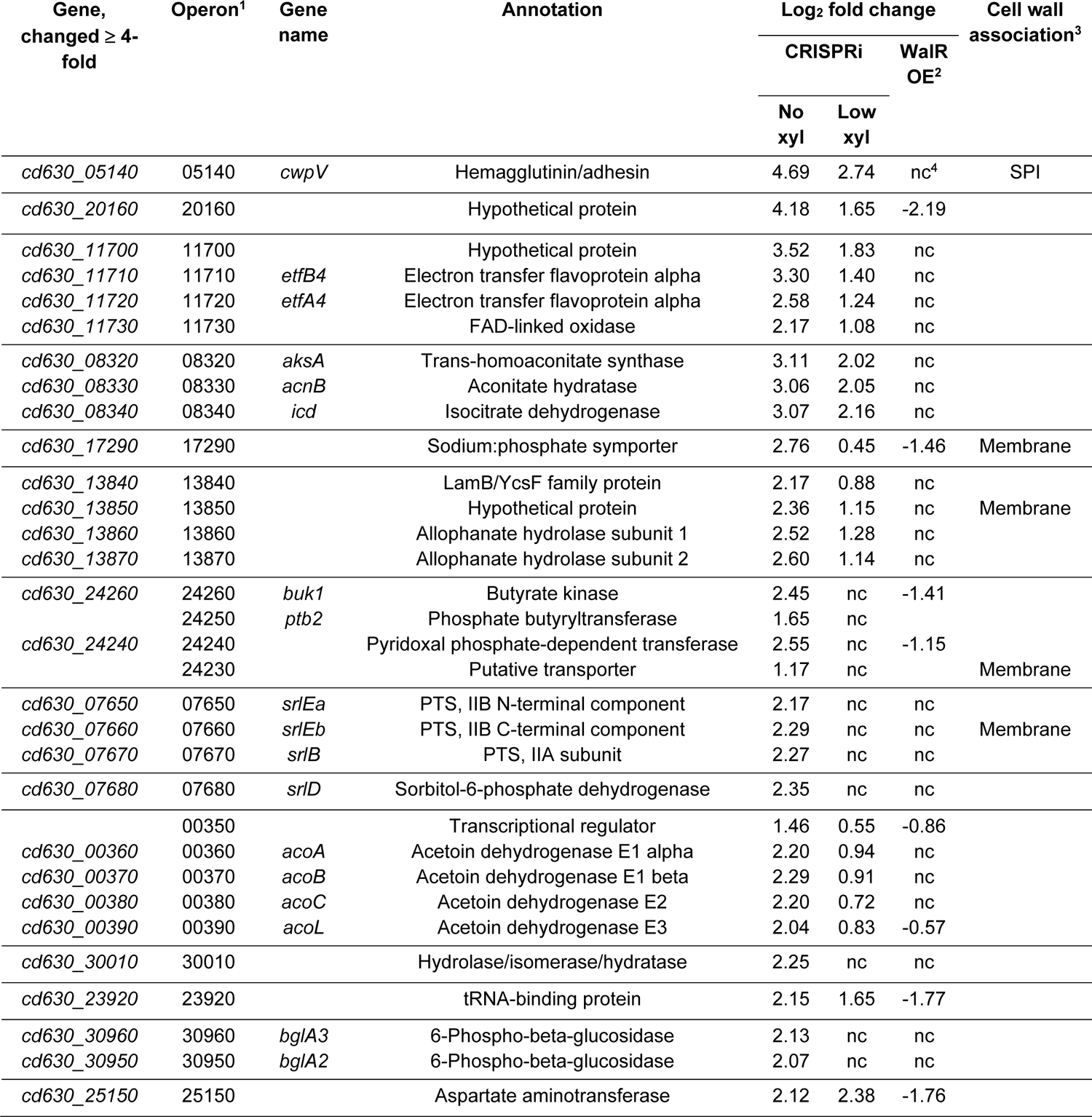
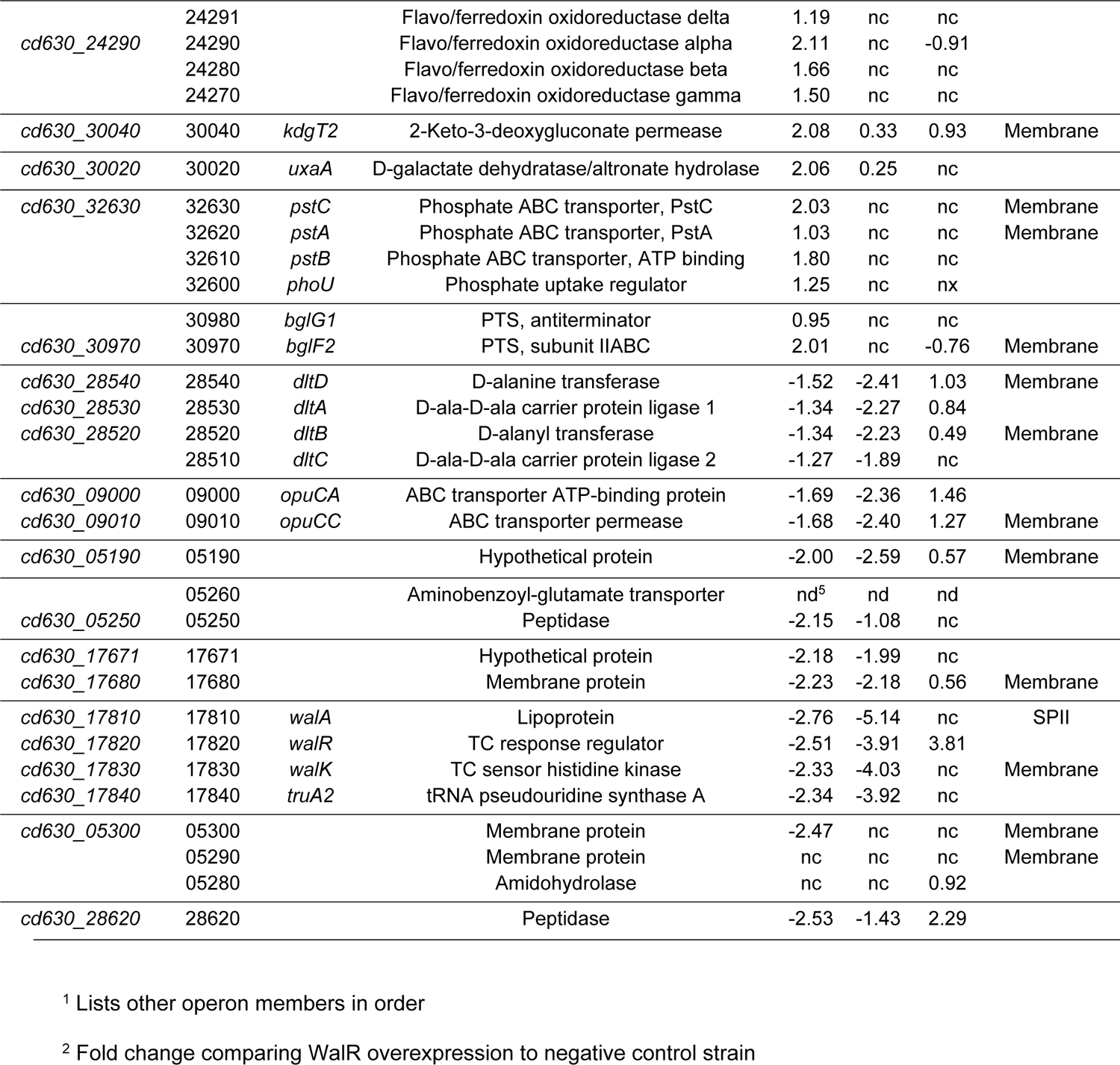
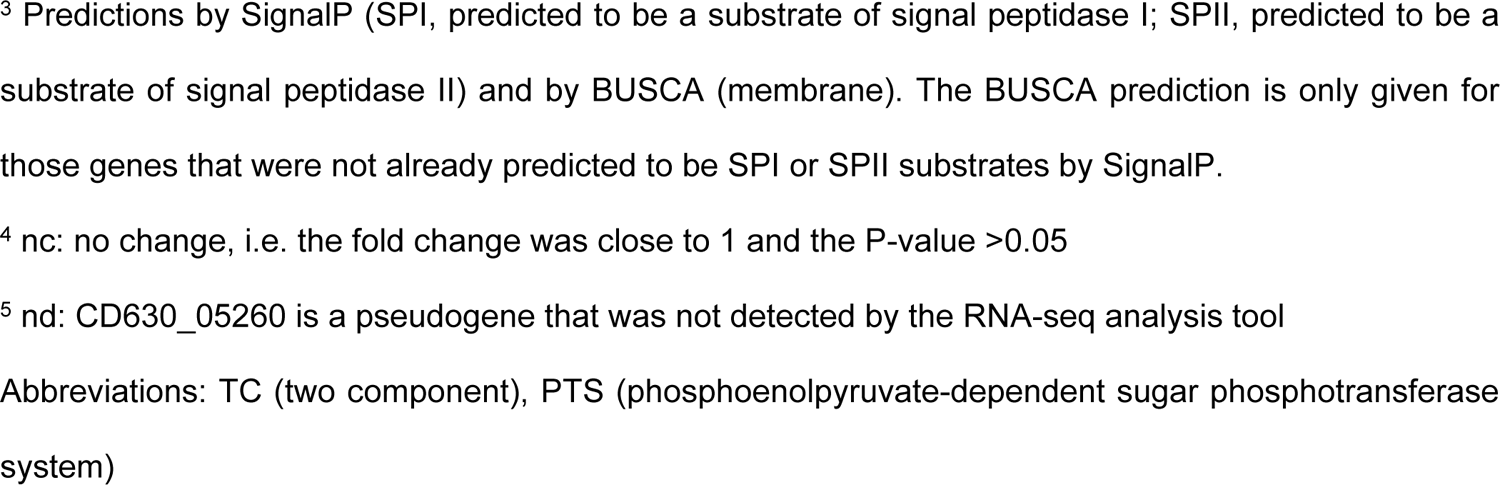
Genes affected by Wal-OFF conditions

The CRISPRi guide targets the first gene in the operon, *walA*, which was repressed 6.8-fold in the absence of xylose and 35-fold with low xylose. As expected, polarity resulted in repression of the downstream genes as well, ∼5-fold with leaky CRISPRi, increasing to ∼15-fold with low xylose. The observed polarity confirms the predicted operon structure, including that a seemingly unrelated (and according to Tn-seq nonessential) gene for a tRNA modification enzyme is indeed part of the operon.

Unexpectedly, the Wal-OFF gene set comprises mostly transporters and genes for various metabolic functions. Not counting the members of the *wal* operon, only one of the genes is predicted to be exported and 15 genes (25%) to be membrane proteins, a percentage comparable to the genome as a whole (Methods). Remarkably, the Wal-OFF regulon does not include a single annotated PG metabolism gene. However, we observed ∼5-fold repression of the *dltDABC* operon, which is responsible for the addition of D-alanine to teichoic acids (49). We identified a predicted WalR binding site immediately upstream of *dltDABC*, suggesting direct regulation by WalR. This putative WalR binding site had previously been recognized as one of two direct repeat sequences in the *dlt* promoter region ((49), Table S4). D-alanylation of teichoic acids affects PG hydrolase activity (50, 51) and vancomycin resistance in some bacteria (52, 53). In *C. difficile*, a *dltD* insertion mutant is sensitized to vancomycin and some antimicrobial peptides, but the effects are <2-fold (52). Only one gene in Table 3 codes for an S-layer protein—the hemagglutinin/adhesin *cwpV*, which was ∼16-fold induced in no xylose. But this gene is subject to phase variation (54), leading us to suspect it may not be a true member of the WalR regulon.

### Comparison of the Wal-ON and the Wal-OFF gene sets reveals little overlap but opposing trends

There are only two proteins common to the Wal-ON and the Wal-OFF gene sets, *cd630_20160* (hypothetical protein) and *cd630_28620* (annotated as a membrane peptidase). However, a large fraction of the 79 Wal-ON genes that met the 4-fold cut-off trend in the opposite direction when evaluated under Wal-OFF conditions, even though the magnitude of change is not as large (Fig. 5A, Table 2). A similar observation was made when genes affected 4-fold or more under Wal-OFF conditions were examined under Wal-ON conditions (Fig. 5B, Table 3).

**Fig. 5.**
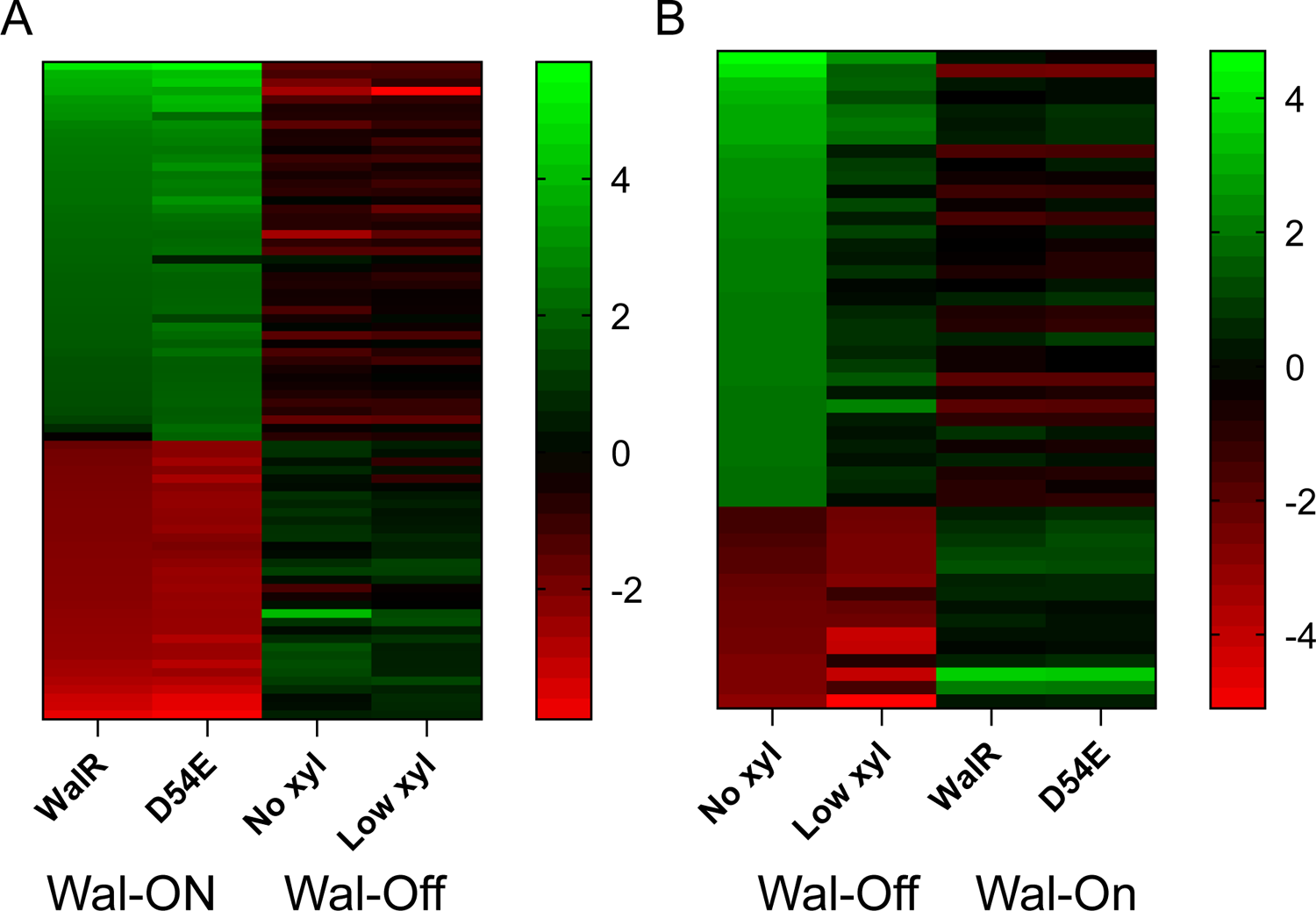
Comparison of transcript changes between Wal-ON and Wal-OFF conditions. (A). Heatmap showing the 79 genes whose transcript abundance changed ≥ 4-fold when *walR* was overexpressed (Wal-ON) in comparison to when the *wal* operon was silenced with CRISPRi (Wal-OFF). (B) Heatmap showing the 49 genes whose transcript abundance changed ≥ 4-fold when the *wal* operon was silenced (Wal-OFF) in comparison to when *walR* was overexpressed (Wal-ON). Green, induced. Red, repressed.

### Select members of the WalR regulon were confirmed by plasmid-based reporter fusions

To confirm results of the RNA-seq analysis, we constructed *rfp* transcriptional fusions (55) to the promoter regions for seven genes from the Wal-ON or Wal-OFF gene sets. Four genes were chosen primarily because they were highly-induced when WalR was overexpressed: *cd630_07390*, *pgdA*, *cd630_07380*, and *cd630_08670* (Table 2). Additional considerations included that *pgdA* has an obvious connection to PG biogenesis (36, 37), while *cd630_08670* has a predicted WalR-binding site (Table S4). Two genes were chosen because they were the most strongly repressed upon CRISPRi knockdown of the *wal* operon and are preceded by a predicted WalR-binding site: *cd630_28620* and *cd630_05300* (Tables 3, S4). In addition, *cd630_28620* is one of only two genes that made the 4-fold cut-off in both Wal-ON and Wal-OFF conditions. The final gene chosen for confirmation with an *rfp* reporter was *dltD*, which incorporates D-alanine into teichoic acids (49) and was modestly downregulated by CRISPRi knockdown of the *wal* operon. *dltD* stood out because it has a predicted WalR-binding site and is one of the few cell envelope-associated genes that was at least 4-fold repressed in Wal-OFF conditions (Tables 3, S4).

Reporter plasmids were constructed by PCR-amplifying ∼300 bp upstream of the start codon for each selected gene. These fragments contain transcriptional start sites for five of the seven genes (Clost-Base database (56)); start sites for the remaining two genes have not been mapped. The PCR products were cloned into an RFP reporter plasmid. The resulting plasmids were conjugated into the appropriate Wal-ON and Wal-OFF strain pairs: 630Δerm versus 630Δerm P*_xyl_::walR* and 630Δerm with sgRNA*-walA1* versus 630Δerm sgRNA-neg, respectively. Reporter strains were grown under the same conditions as used for the RNA-seq experiments, except that the Wal-OFF condition was limited to growth in the absence of xylose, i.e., assaying the effect of leaky dCas9 expression on the reporter gene. Harvested cells were fixed and transferred to aerobic conditions to allow RFP to mature. Fluorescence was quantified by flow cytometry.

We found that all RFP reporters responded in the expected direction when *walR* expression was manipulated, thus confirming the RNA-seq results by an orthogonal method (Fig. 6). However, the fold changes were uniformly about 4-fold greater by RNA-seq than by RFP fluorescence. The reasons for this difference are not known. As an aside, having RNA-seq and RFP fluorescence data for the same genes enabled us to show these are correlated, i.e., baseline RFP fluorescence was higher for genes that returned more reads in RNA-seq experiments (Fig. S5A). This correlation was expected and further indicates our RNA-seq data are robust.

**Fig. 6.**
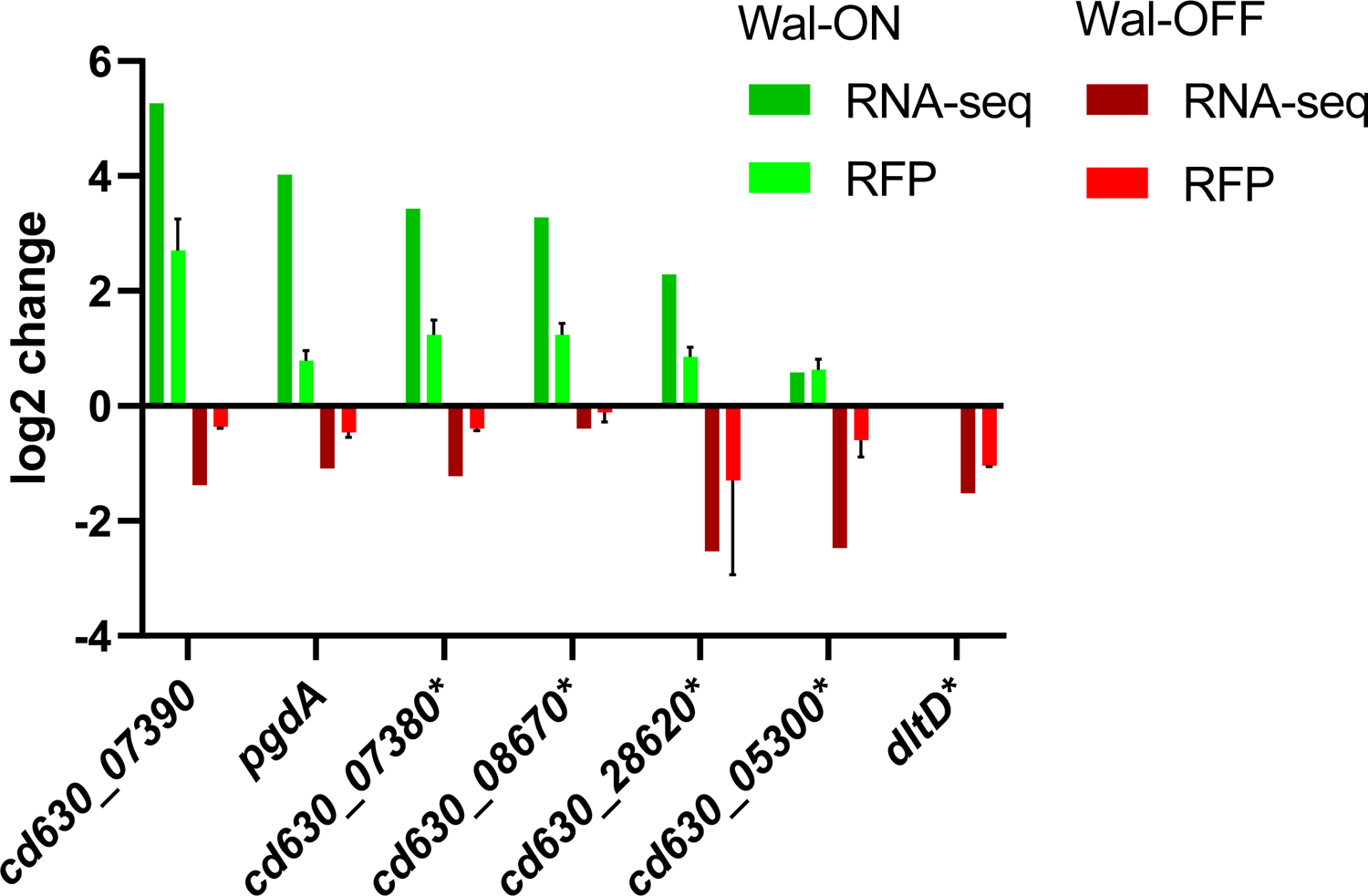
Confirmation of RNA-seq results with RFP reporter fusions to select genes. Cells harboring plasmids with transcriptional fusions of the red fluorescent protein mCherryOpt to the indicated promoters were grown as per RNA-seq conditions, either Wal-ON (green) or Wal-OFF (red). Red fluorescence was measured by flow cytometry and is graphed as the mean log2-fold change from two independent experiments with two technical replicates. Error bars represent the SD of all four measurements. For comparison, the mean log2-fold change of the same genes as determined by RNA-seq is also shown. The *dltD* reporter is graphed only under Wal-OFF conditions because this fusion was induced by xylose even in the absence of P*_xyl_*::*walR*. Genes with asterisks denote fusions that were constructed to the corresponding promoter regions from strain R20291.

### Altering the expression level of select regulon genes does not induce the *wal* reporter

In *B. subtilis*, the WalR regulon can be induced or repressed by manipulating expression of *walR*-regulated genes involved in PG metabolism (11, 57). We therefore asked whether manipulating expression of *wal* regulon genes in *C. difficile* would induce or repress expression of a chromosomal P*_cd630_07390_*::*rfp* reporter. The reporter was chosen because this promoter was highly induced in Wal-ON conditions as determined by RNA-seq (38-fold) or a plasmid-based RFP reporter fusion (7-fold). The chromosomal P*_cd630_07390_*::*rfp* reporter was validated by confirming ∼10-fold increased red fluorescence upon overexpression of *walR* from a P*_xyl_* plasmid (Fig. S5B). Next, we introduced a panel of plasmids that allowed us to directly target 13 WalR regulon genes by overexpression and/or CRISPRi. These 13 genes were chosen for various reasons including large fold-changes in expression, relevance to cell wall biogenesis, and the presence of WalR binding sites (Table S3). Of note, the gene set included a putative cell wall amidase and the predicted PG hydrolase with a NlpC/P60 domain. Unfortunately, neither overexpression nor CRISPRi knockdown of the selected *wal* regulon genes altered expression of the P*_cd630_07390_*::*rfp* reporter. Further work will be needed to determine whether these negative results reflect technical difficulties or more fundamental differences in Wal signaling between *B. subtilis* and *C. difficile*.

### Perturbation of select Wal regulon genes individually does not lead to any Wal-associated phenotypic defects

As noted above, one motivation for determining the Wal regulon was to identify genes that contribute to Wal phenotypes. Recall that induction of P*_xyl_*::*walR* decreases viability, causes cells to become phase bright and reduces autolysis in buffer containing 0.01% Triton X-100 (Fig. 3). Two of the most highly induced genes in Wal-ON conditions encode the hypothetical proteins CD630_07380 and CD630_07390. However, overproduction of these proteins either individually or from P*_xyl_* plasmids did not result in any obvious phenotypic changes (Fig. 7). Note, because these genes are transcribed divergently (35), we needed to reverse the direction of one gene to express them together and decided to clone both possible arrangements: *cd630_07380*-*cd630_07390* and *cd630_07390*-*cd630_07380*. Likewise, a CRISPRi plasmid with an sgRNA that targets *cd630_27680* had no effects on growth, morphology or lysis (Fig. 7); this gene encodes PG hydrolase with two predicted WalR-binding sites and is among the most strongly repressed genes in Wal-ON conditions.

**Fig. 7.**
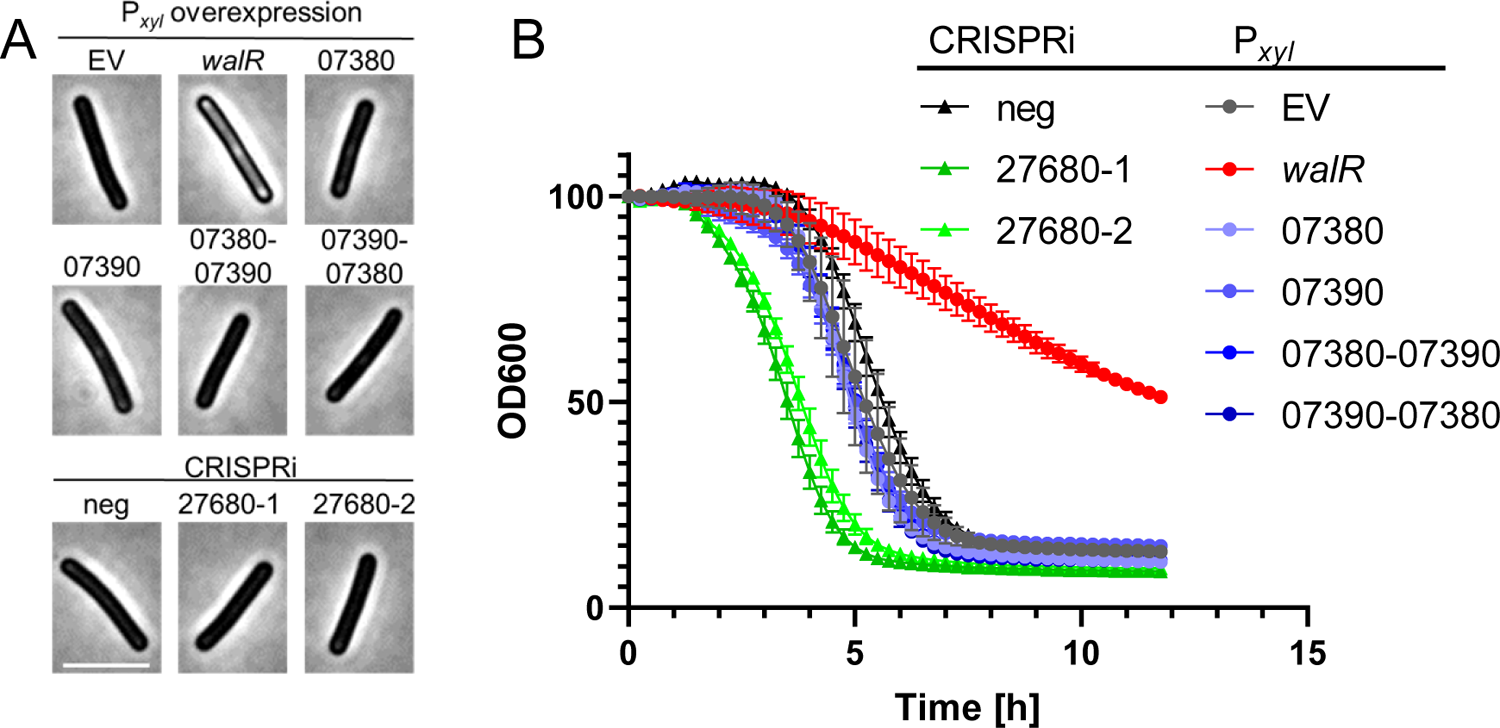
Perturbation of individual Wal-ON regulon genes does not replicate the Wal-ON phenotype. Overnight cultures of 630Δerm harboring overexpression plasmids were subcultured into TY-Thi to an OD_600_ of 0.03, grown to OD_600_ = 0.2 and induced with 3% xylose for 3h. Overnight cultures of 630Δerm harboring CRISPRi plasmids were subcultured into TY-Thi containing 1% xylose to an OD_600_ of 0.03 and grown to OD_600_ ∼0.8. Each culture was then (A) examined by microscopy (bar 5 µm) and (B) tested in the lysis assay. Induction of *walR* is the only condition that achieved phase bright cells and slowed lysis. The plasmids used were: pBZ101 (EV), pCE691 (P*_xyl_*::*walR*), pIA112 (P*_xyl_*::*cd630_07380*), pIA113 (P*_xyl_*::*cd630_07390*), pIA114 (P*_xyl_*::*cd630_07380-07390*), pIA115 (P*_xy_*_l_::*cd630_07390-07380*), pIA34 (sgRNA-neg), pCE744 (*sgRNA-cd630_27680-1*), and pCE745 (*sgRNA-cd630_27680-2*).

CRISPRi silencing of the *wal* operon is lethal, and the terminal phenotypes include lysis and loss of rod shape as reflected by an abundance of curved cells (Fig. 2). However, none of these phenotypes were observed when we used a P*_xyl_*::*cwpV* plasmid to overexpress the most highly-induced gene from the Wal-OFF RNA-seq data set, which we tested even though *cwpV* is subject to phase-variation and might not be part of the WalR regulon. We also observed an effect when we used CRISPRi to knock down expression of *dltD*, which was among the more strongly repressed Wal-OFF genes and is preceded by a putative WalR binding site (Fig. 8). While not an exhaustive evaluation of all regulon members, our findings suggest the phenotypic defects observed upon perturbation of the Wal system are not due to changes in expression of any single gene, but cumulative in nature much like what has been observed in *B. subtilis* and *S. aureus* (8–10, 12).

**Fig. 8.**
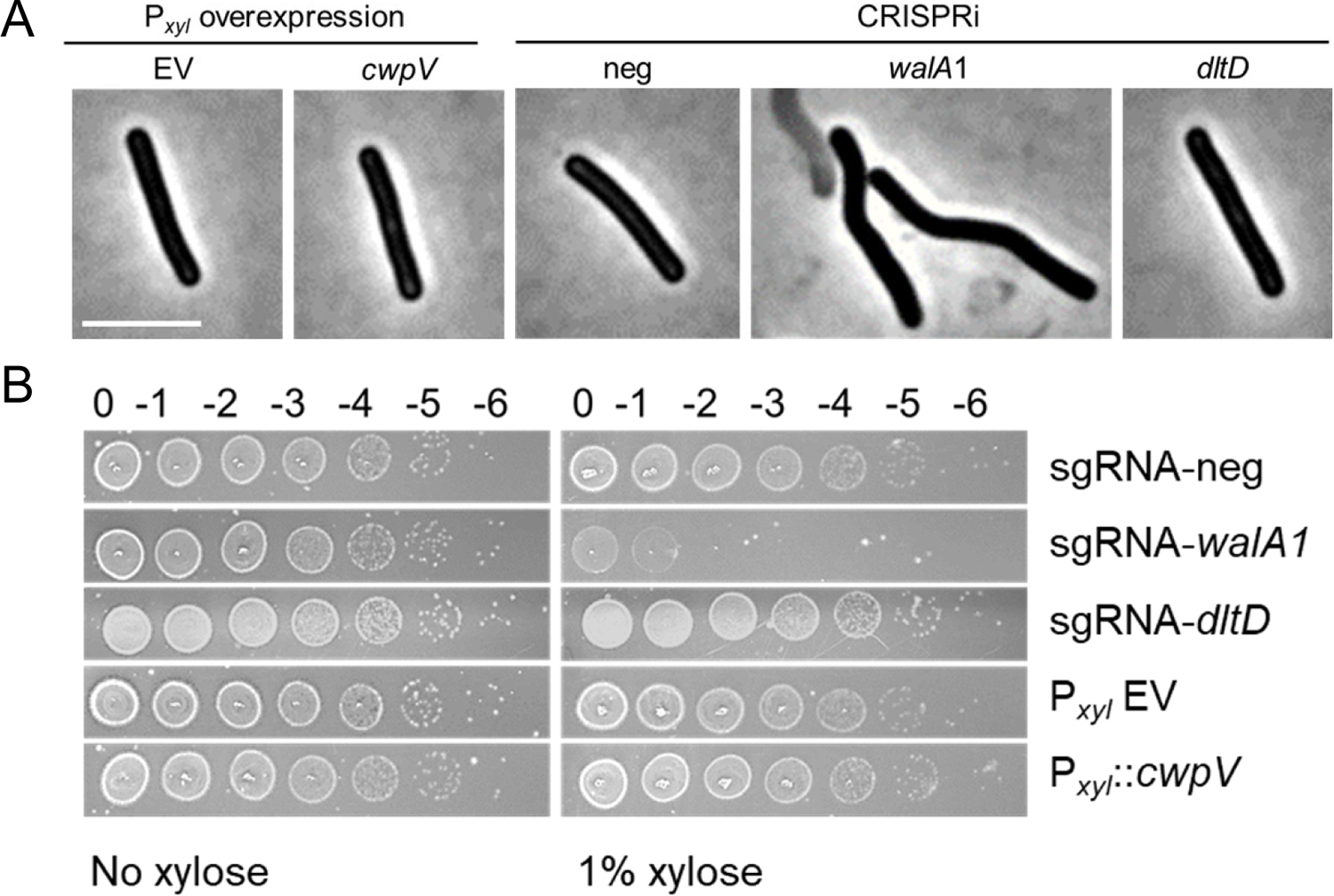
Perturbation of individual Wal-OFF regulon genes does not replicate the Wal-OFF phenotype. (A) Phase contrast microscopy of strains harboring overexpression plasmids or CRISPRi plasmids after induction with xylose as described in the legend to Fig. 7. Bar = 5 µm. (B) Viability assay. Overnight cultures were serially diluted and spotted onto TY plates with or without 1% xylose. Plates were photographed after incubation overnight. CRISRPi knockdown of the *wal* operon was the only condition that resulted in curved cells, lysis or a viability defect. The plasmids used were pBZ101 (EV), pCE791 (P*_xyl_*::*cwpV*), pIA34 (sgRNA-neg), pIA50 (sgRNA-*walA1*), and pCE738 (sgRNA-*dltD*).

## Discussion

In *C. difficile* the WalRK TCS is predicted to reside in a four gene operon: *walA-walR-walK-truA2*. We altered expression of the WalRK regulon by complementary approaches: depleting *C. difficile* of WalRK by CRISPRi knockdown of the entire operon, and overproduction of WalR under P*_xyl_* control. RNA-seq revealed the *wal* operon is not autoregulated in *C. difficile*, as evidenced by a lack of induction when WalR (or WalR^D54E^) was ectopically expressed from a P*_xyl_* promoter. Wal operons are not autoregulated in other Firmicutes either. RNA-seq also confirmed the genes constitute an operon, because CRISPRi targeting *walA* resulted in roughly equivalent knockdown of the three downstream genes. Further studies will be needed to establish the specific role of each gene in WalRK signaling. We suspect *walA* is a bona fide part of the Wal system because all *wal* operons studied to date include accessory genes for membrane proteins that modulate WalK signaling (8, 10). But a rationale for why a non-essential tRNA modification enzyme, *truA2,* should have come to reside in the *wal* operon is not obvious. This might be the result of evolutionary happenstance.

As expected, extensive knockdown of the *wal* operon with CRISPRi or strong overproduction of WalR from a P*_xyl_* plasmid were both lethal. Phenotypic defects depended on how *walRK* expression was manipulated, but included elongation, loss of rod shape (curved or wavy cells), phase bright cells, enhanced or reduced autolysis, and altered sensitivity to antibiotics that target PG synthesis. Similar phenotypes have been observed upon manipulating Wal signaling in other organisms (2, 6, 7, 9, 10, 12, 17, 18). But there are some intriguing differences. In *B. subtilis*, where phosphorylated WalR promotes elongation, artificial upregulation of the Wal system causes cells to become elongated while artificial downregulation causes cells to become short (10, 18). In *C. difficile*, however, overexpression of WalR from P*_xyl_* had no obvious effect on cell length while CRISPRi knockdown of the *wal* operon led to elongation.

Vancomycin resistance presents another example of an inverse phenotype. In *S. aureus* down regulating *walRK* expression increases resistance to vancomycin while up regulation of *walRK* decreases vancomycin resistance (58, 59). Vancomycin-intermediate *S. aureus* (VISA) strains, both laboratory derived (60) or isolated in the clinic (61), often have mutations in *walR* or *walK* that down regulate the system (62). In contrast, we found that partial CRISPRi knockdown of the *wal* operon in *C. difficile* had the opposite effect, namely, a modest increase in sensitivity to vancomycin. Vancomycin is a front line treatment for *C. difficile* infections, so our findings suggest a small molecule that interferes with Wal signaling might enhance the efficacy of vancomycin therapy.

RNA-seq identified over 150 genes whose expression responds to manipulation of Wal signaling in *C. difficile*. Although this gene set is diverse, there are themes: many cell envelope proteins, many hypotheticals, and about 10 proteins with various connections to PG metabolism. There are also 7 genes for S-layer proteins, consistent with the need to coordinate S-layer assembly and remodeling with growth and turnover of the PG sacculus. The PG hydrolases are of particular interest because they are prominent members of the WalR regulon in other Firmicutes (5, 12–16) and ectopic expression of one or more of these enzymes can render *walRK* no longer essential regulators (6, 17, 18). In *C. difficile*, Wal-ON conditions induced two putative cell wall amidases (*cwp16*, *cwp17*) and repressed one potential D,L-endopeptidase (*cd630_27680*). Direct control of *cd630_27680* by WalR is suggested by the presence of two matches to the WalR binding site consensus sequence about 100 nucleotides upstream of the start codon (Table S4). The importance of these genes for any Wal-related phenotypes is unclear, however, because overexpression and CRISPRi knockdown experiments did not result in any phenotypic changes or alter expression of an RFP reporter that is induced under Wal-ON conditions. In addition to the putative hydrolases, several genes in the WalR regulon may play a role in *controlling* PG hydrolase activity. Deacetylation of PG N-acetylglucosamine (GlcNAc) (63), mDAP amidation (39), and D-alanylation of teichoic acids (32, 50, 51, 64) have all been shown to affect hydrolase activity in other organisms.

*S. pneumoniae* is so far the only organism in which a single essential gene, the PG hydrolase *pcsB*, has been identified to be responsible for the essentiality of the *wal* system (6, 7). In *B. subtilis*, essentiality of the Wal system arises from the cumulative impact of aberrant expression of multiple genes involved in turnover of PG (5, 10, 11). Wal essentiality is also polygenic in nature in *S. aureus* (8). This seems to be the case in *C. difficile* as well. Although CRISPRi silencing of the *wal* operon leads to cell lysis, the Wal-OFF gene set (Table 3) does not include any genes determined to be essential by transposon mutagenesis (27). In contrast, the Wal-ON regulon (Table 2) includes 3 essential genes. Two of these are induced by WalR, DNA ligase (*ligA, cd630_33090*) and a putative Calcium-chelating protein (*cd630_26810*); one is repressed, the PG precursor synthesis protein MurE (*cd630_26640*). It is possible that repression of *murE* contributes to death during overexpression of WalR, but not during CRISPRi knockdown of the *wal* operon, because *murE* expression did not change under that condition. Overall, our findings suggest essentiality of WalRK is driven by a deleterious imbalance of multiple processes. (As an aside, it is worth noting that repression of *murE* under Wal-ON conditions is not what one would expect if WalR promotes increased PG synthesis, as in *B. subtilis* (11, 57)).

Here we have provided an initial delineation of Wal phenotypes and the WalR regulon in *C. difficile*. Our findings raise a number of exciting questions. Is the protein WalA involved in WalRK signaling, and if so, how? What signals are sensed by WalK? Of note, owing to differences in crosslinking (23), the PG hydrolysis products implicated in regulating WalK activity in *B. subtilis* (11) are not very abundant in *C. difficile*. Moreover, *C. difficile* WalK is missing an intracellular PAS domain found in most other WalK proteins, suggesting *C. difficile* WalK senses different signals and/or transmits those signals differently. Another open question is which WalR regulon genes are controlled directly by WalR. It will also be important to determine the functions of the WalR regulated genes, especially the many hypothetical proteins. Answers to these questions are likely to provide novel insights into cell wall biogenesis and might point the way towards improved therapies against this important pathogen.

## Methods

### Strains, media, and growth conditions

Bacterial strains are listed in Table 4. *C. difficile* strains used in this study were derived from either 630Δerm or R20291, both of which have been sequenced. *C. difficile* was routinely grown in tryptone-yeast extract (TY) medium, supplemented as needed with thiamphenicol at 10 μg/ml (TY-Thi). TY medium consisted of 3% tryptone, 2% yeast extract, and 2% agar (for plates). Brian heart infusion (BHI) media was prepared per manufacturer’s (DIFCO) instructions. *C. difficile* strains were maintained at 37°C in an anaerobic chamber (Coy Laboratory Products) in an atmosphere of 10% H_2_, 5% CO_2_, and 85% N_2_. *Escherichia coli* strains were grown in LB medium at 37°C with chloramphenicol at 10 μg/ml or ampicillin at 100 μg/ml as needed. LB medium contained 1% tryptone, 0.5% yeast extract, 0.5% NaCl, and 1.5% agar (for plates). OD_600_ measurements were made with the WPA Biowave CO8000 tube reader in the anaerobic chamber.

**Table 4.** Strains used in this study

| Strain | Genotype and/or description | Source or reference |
| --- | --- | --- |
| *E. coli* |  |  |
| OmniMAX-2 T1<sup>R</sup> | F' [*proAB*<sup>+</sup> *lacI*<sup>q</sup> *lacZ*ΔM15 Tn10(Tet<sup>R</sup>) Δ(*ccdAB*)] *mcrA* Δ(*mrr-hsdRMS-mcrBC*) φ80(*lacZ*)ΔM15 Δ(*lacZYA-argF*) U169 *endA1 recA1 supE44 thi-1 gyrA96 relA1 tonA panD* | Invitrogen |
| HB101/pRK24 | F– *mcrB mrr hsdS20*(r<sub>B</sub><sup>–</sup> m<sub>B</sub><sup>–</sup>) *recA13 leuB6 ara-14 proA2 lacY1 galK2 xyl-5 mtl-1 rpsL20* | (66, 67) |
| *C. difficile* |  |  |
| R20291 | Wild-type *C. difficile* strain from UK outbreak (ribotype 027) |  |
| 630Δ*erm* | Spontaneous erythromycin-sensitive derivative of strain 630 (ribotype 012) | (76) |
| CRG1496 | 630Δ*erm*Δ*pyrE* | (29) |
| UM275 | 630Δ*erm* with P<sub>veg</sub>::*rfp* downstream of *pyrE* | (26) |
| UM554 | 630Δ*erm* with P<sub>xyl</sub>::*dCas9* P<sub>gdh</sub>::*sgRNA*-neg downstream of *pyrE* | This study |
| UM555 | 630Δ*erm* with P<sub>xyl</sub>::*dCas9* P<sub>gdh</sub>::*sgRNA-walA-1* downstream of *pyrE* | This study |
| UM556 | 630Δ*erm* with P<sub>xyl</sub>::*dCas9* P<sub>gdh</sub>::*sgRNA-walA-2* downstream of *pyrE* | This study |
| UM626 | 630Δ*erm* with P<sub>xyl</sub>::*walR* downstream of *pyrE* | This study |
| UM628 | 630Δ*erm* with P<sub>xyl</sub>::*walR*<sup>D54E</sup> downstream of *pyrE* | This study |
| UM926 | 630Δ*erm* with P<sub>cd630_0739</sub>::*mCherryOpt* downstream of *pyrE* | This study |

### Plasmid and strain construction

All plasmids are listed in Table 5; an expanded version of this table which includes additional information relevant to plasmid assembly is provided in Table S2A in the supplemental material. Plasmids were constructed by isothermal assembly (65) using reagents from New England Biolabs (Ipswich, MA). Regions of plasmids constructed using PCR were verified by DNA sequencing. The oligonucleotide primers used in this work were synthesized by Integrated DNA Technologies (Coralville, IA) and are listed in Table S2B. All plasmids were propagated using OmniMax 2-T1R as the cloning host, transformed into HB101/pRK24 (66, 67), and then introduced into *C. difficile* strains by conjugation. Chromosomal fusions at the *pyrE* locus in 630Δerm were constructed by allelic exchange (29) using *C. difficile* CRG1496 (630Δ*erm* Δ*pyrE*) as a *pyrE*-deficient recipient. The allelic exchange restored a functional *pyrE* gene.

**Table 5.** Plasmids used in this study

| Plasmid | Relevant features | Reference or comment |
| --- | --- | --- |
| pAP114 | *P<sub>xyl</sub>::mCherryOpt catP* | (26) |
| pBZ101 | *P<sub>xyl</sub>* empty vector | This study |
| pCE636 | *P<sub>dltD</sub>::mCherryOpt catP* | This study |
| pCE738 | *P<sub>xyl</sub>::dCas9-opt P<sub>gdh</sub>::sgRNA-dltD-1 catP* | (37) |
| pCE691 | *P<sub>xyl</sub>::walR catP* | This study |
| pCE738 | *P<sub>xyl</sub>::dCas9-opt P<sub>gdh</sub>::sgRNA-dltD-1 catP* | (37) |
| pCE741 | *P<sub>xyl</sub>::cdr2656* | This study |
| pCE744 | *P<sub>xyl</sub>::dCas9-opt P<sub>gdh</sub>::sgRNA-cd630_27680-1 catP* | This study |
| pCE745 | *P<sub>xyl</sub>::dCas9-opt P<sub>gdh</sub>::sgRNA-cd630_27680-2 catP* | This study |
| pCE789 | *P<sub>xyl</sub>::dCas9-opt P<sub>gdh</sub>::cwpV-1 catP* | This study |
| pCE791 | *P<sub>xyl</sub>::cwpV* | This study |
| pDSW1728 | *P<sub>tet</sub>::mCherryOpt catP* | (55) |
| pDSW2037 | *P<sub>xyl</sub>* in integration vector pMTL-YN1C *catP* | This study |
| pDSW2053 | *P<sub>xyl</sub>::dCas9opt-P<sub>gdh</sub>::sgRNA-neg* in pMTL-YN1C *catP* | This study |
| pDSW2055 | *P<sub>xyl</sub>::dCas9opt-P<sub>gdh</sub>::sgRNA-cd630_17810-1* in pMTL-YN1C *catP* | This study |
| pDSW2057 | *P<sub>xyl</sub>::dCas9opt-P<sub>gdh</sub>::sgRNA-cd630_17810-2* in pMTL-YN1C *catP* | This study |
| pIA33 | *P<sub>xyl</sub>::dCas9-opt P<sub>gdh</sub>::sgRNA-rfp catP* | (26) |
| pIA34 | *P<sub>xyl</sub>::dCas9-opt P<sub>gdh</sub>::sgRNA-neg catP* | (26) |
| pIA50 | *P<sub>xyl</sub>::dCas9-opt P<sub>gdh</sub>::sgRNA-walA-1 catP* | This study |
| pIA51 | *P<sub>xyl</sub>::dCas9-opt P<sub>gdh</sub>::sgRNA-walA-2 catP* | This study |
| pIA75 | *P<sub>xyl</sub>::walR* in pMTL-YN1C *catP* | This study |
| pIA76 | *P<sub>xyl</sub>::walR<sup>D54E</sup>* in pMTL-YN1C *catP* | This study |
| pIA79 | *P<sub>xyl</sub>::dCas9-opt P<sub>gdh</sub>::cd630_25040-1 catP* | This study |
| pIA80 | *P<sub>xyl</sub>::dCas9-opt P<sub>gdh</sub>::cd630_25040-2 catP* | This study |
| pIA81 | *P<sub>xyl</sub>::dCas9-opt P<sub>gdh</sub>::cd630_36010-1 catP* | This study |
| pIA93 | *P<sub>cdr_0665</sub>::mCherryOpt catP* | This study; *cdr_0665* is ortholog of *cd630_07380* |
| pIA95 | *P<sub>cdr_0796</sub>::mCherryOpt catP* | This study; *cdr_0796* is ortholog of *cd630_08670* |
| pIA97 | *P<sub>cdr_2753</sub>::mCherryOpt catP* | This study; *cdr_2753* is ortholog of *cd630_28620* |
| pIA98 | *P<sub>cdr_0455</sub>::mCherryOpt catP* | This study; *cdr_0455* is ortholog of *cd630_53000* |
| pIA100 | *P<sub>cd630_0739</sub>::mCherryOpt catP* | This study |
| pIA101 | *P<sub>cd630_0739</sub>::mCherryOpt* in pMTL-YN1C *catP* | This study |
| pIA102 | *P<sub>xyl</sub>::dCas9-opt P<sub>gdh</sub>::cd630_05490 catP* | This study |
| pIA103 | *P<sub>xyl</sub>::dCas9-opt P<sub>gdh</sub>::cd630_27940 catP* | This study |
| pIA104 | *P<sub>xyl</sub>::dCas9-opt P<sub>gdh</sub>::cd630_27860 catP* | This study |
| pIA105 | *P<sub>xyl</sub>::dCas9-opt P<sub>gdh</sub>::cd630_27910 catP* | This study |
| pIA106 | *P<sub>xyl</sub>::dCas9-opt P<sub>gdh</sub>::cd630_10360 catP* | This study |
| pIA107 | *P<sub>pgdA</sub>::mCherryOpt catP* | This study |
| pIA108 | *P<sub>xyl</sub>::dCas9-opt P<sub>gdh</sub>::cd630_07380-1 catP* | This study |
| pIA109 | *P<sub>xyl</sub>::dCas9-opt P<sub>gdh</sub>::cd630_07380-2 catP* | This study |
| pIA110 | *P<sub>xyl</sub>::dCas9-opt P<sub>gdh</sub>::cd630_07390-1 catP* | This study |
| pIA111 | *P<sub>xyl</sub>::dCas9-opt P<sub>gdh</sub>::cd630_07390-2 catP* | This study |
| pIA112 | *P<sub>xyl</sub>::cd630_0738* | This study |
| pIA113 | *P<sub>xyl</sub>::cd630_0739* | This study |
| pIA114 | *P<sub>xyl</sub>::cd630_0738-cd630_0739* | This study |
| pIA115 | *P<sub>xyl</sub>::cd630_0739-cd630_0738* | This study |
| pMTL-YN1C | *E. coli-C. difficile* shuttle vector for inserting genes into *C. difficile* chromosome while restoring *pyrE*; *colE1 RP4oriT-TraJ CB102ori-repH'* *catP* | (29) |

### Conjugation into *C. difficile*

Our experiments required conjugating plasmids into *C. difficile* strains R20291 and 630Δerm. In the case of R20291 we encountered problems with low conjugation efficiencies even with heat shock as described (68). After testing several modifications of the procedure, we found that performing conjugations on filters reliably increased efficiency more than 10-fold (Fig. S6). We now routinely use filters for all our conjugations. Briefly, the *E. coli* donor strain (an HB101/pRK24 derivative harboring the cargo plasmid) was grown overnight in LB Amp^100^ Cam^10^. The donor strain was collected gently by centrifugation of a 0.5 ml aliquot at 5000 x g for 1 min, then washed with 1 ml TY and pelleted again. The washed cell pellet was moved into the anaerobic chamber. R20291 was prepared for conjugation by a heat shock step (68). For this, a 200 µl aliquot of the overnight culture of the R20291 recipient was transferred to a 1.5 ml microfuge tube and incubated at 48°C for 5 min in a Fisherbrand Dry bath (with water-filled wells). Then 100 µl of heat-shocked R20291 was used to take up the pellet of *E. coli* donor cells. The strain mixture was then pipetted on to a 25 mm diameter, 0.45 μm pore size Millipore mixed cellulose filter (HAWP02500) placed on brain heart infusion (BHI) plates. After incubation for 24 h at 37°C, cells were flushed from the membrane with 500 μL TY and 200-500 μL of the resulting bacterial slurry was plated on TY amended with thiamphenicol (10 μg/ml), kanamycin (50 μg/ml) and cefoxitin (8 μg/ml) to select for exconjugants. Conjugations into 630Δerm were done identically except the heat shock step was omitted.

### Viability assay

The effect of CRISPRi silencing on plating efficiency was evaluated by making a 10-fold serial dilution of a culture grown overnight in TY Thi10 and spotting 5 μl of each dilution on TY Thi10 agar with and without 1% xylose. Plates were photographed after overnight incubation (∼18 h).

### Microscopy

Cells were immobilized using thin agarose pads (1%). Phase-contrast and fluorescent micrographs were recorded on an Olympus BX60 microscope equipped with a 100× UPlanApo objective (numerical aperture, 1.35). Micrographs were captured with a Hamamatsu Orca Flash 4.0 V2+ complementary metal oxide semiconductor (CMOS) camera. The image analysis tool MicrobeJ (69) was used to measure cell length and sinuosity, which is the length of the cell axis divided by the distance between the poles. A perfectly straight rod has a sinuosity value of 1, while a curved or wavy cell has a larger value. We classified cells as curved if their sinuosity was ≥1.03 (see Fig. S2 for examples). Viability of cells was assessed with LIVE/DEAD stain (Molecular Probes L7012). A 1 mL culture sample was pelleted, washed with PBS and resuspended in 100 μL PBS with 5 μM Syto 9 and 30 μM propidium iodide. Cells were incubated with the dyes for 15 min, then removed from the anaerobic chamber and immediately imaged by microscopy. For propidium iodide red fluorescence we used filter set 41004 (Chroma Technology) with a 538 to 582-nm excitation filter, 595-nm dichroic mirror (long pass), and a 582 to 682-nm emission filter. Syto 9 green fluorescence was captured with filter set 41017 (Chroma Technology Corp) with a 450- to 490-nm excitation filter, a 495-nm dichroic mirror (long pass), and a 500- to 550-nm emission filter.

### Fixation protocol

A 500-μl aliquot of cells in growth medium was added directly to a microcentrifuge tube containing 120 μl of a 5x fixation cocktail: 100 μl of a 16% (wt/vol) paraformaldehyde aqueous solution (Alfa Aesar, Ward Hill, MA) and 20 μl of 1 M NaPO_4_ buffer (pH 7.4). The sample was mixed, incubated in the anaerobic chamber at 37°C for 30 min, then on ice 30 min, and removed from the chamber. The fixed cells were washed twice with 1 mL phosphate-buffered saline (PBS), resuspended in 50 μl of PBS, and left in the dark for 18 h to allow for chromophore maturation.

### MIC determination

Antibiotic sensitivity was determined in 96 well plates. A twofold dilution series of select antibiotics was prepared in 50 μl TY medium. Wells were then inoculated with 50 μl of a diluted culture suspension (10^6^ CFU/ml; calculated OD_600_ ∼0.005). Plates were imaged and evaluated after 17h at 37°C.

### Lysis assay

Cell cultures (1 mL) were removed from the anaerobic chamber, pelleted and resuspended in 700 μL 0.01% Triton X-100 in 50 mM NaPO_4_ buffer (pH 7.4). Of this 700 μL, three 200 μL replicates were pipetted into wells of a clear, flat bottom 96-well plate. The turbidity was measured at 600 nm every 15 min for 10 h in a plate reader (Tecan Infinite M200 Pro).

### Flow cytometry

Cells were analyzed at the Flow Cytometry Facility at the University of Iowa using the Becton, Dickinson LSR II instrument with a 561-nm laser, a 610/20-nm-band-pass filter, and a 600 LP dichroic filter. Data were analyzed using BD FACSDiva software.

### Culture growth for RNA-seq samples

To achieve robust, quality data all experiments were performed on three different days for biological replicates. Cultures were grown in 100 mL TY as follows. Wal-ON: to characterize the effect of overexpressing *walR*, strains were subcultured to OD_600_ ∼0.05, grown to 0.2, induced with xylose to 3% and grown for an additional 3h (Fig. S4A). Wal-OFF: to characterize the effect of partial knockdown of the *wal* operon under leaky CRISPRi conditions, strains were subcultured to OD_600_ ∼0.03 and grown to ∼0.7 before harvesting (Fig. S4C). To test the effect of stronger knockdown of the *wal* operon, growth conditions were identified that allowed exposing cells to moderate levels of xylose (0.1%) without a significant loss of biomass. Strains were subcultured to OD_600_ ∼0.03, grown to 0.3, diluted back to ∼0.1 and induced with xylose to a final concentration of 0.1% and harvested after an additional 2.5 h of growth (Fig. S4D). From each condition, 25 mL culture was fixed by rapidly mixing with 25 mL ice-cold 1:1 ethanol:acetone that had been brought into the anaerobic chamber on dry ice. Fixed cells were stored at −80°C until further workup.

### RNA isolation and RNA-Seq

Fixed culture samples were thawed, pelleted for 10 min at 8000 x g, 4°C and pellets washed once with 500 μL 1% β-mercaptoethanol (β-ME) in water. Subsequent steps are an adaptation of the Qiagen RNeasy Mini Kit procedure. The washed pellet was resuspended in 1 mL of the Qiagen-provided RTL buffer amended with 1% β-ME and transferred to a 2 mL screwcap tube filled with glass beads (0.1 mm diameter) to a height of ∼2-3 mm. Cells were lysed with the FastPrep-24 homogenizer with two cycles at 6m/s for 45 s, resting on ice for 3 min between cycles. The homogenized material was pelleted at 14,000 x g for 15 min at 4°C. The supernatant was removed, adjusted to 900 μL with RTL/β-ME, mixed with 500 μL ethanol and loaded onto the Qiagen RNeasy spin column. Subsequent steps followed the manufacturer’s protocol. The RNA was eluted with 50 μL water and DNase treated twice (Ambion Turbo DNA-free). Absence of DNA was evaluated by amplifying a 364 bp region of *mldA* with Taq DNA polymerase for 30 cycles. The RNA integrity number (RIN) was determined with an Agilent Bioanalyzer at the Iowa Institute for Human Genetics, Genomics Division. All samples had a RIN of 9.4 or higher. Typical yields from 25 mL culture were 8-20 μg RNA at 200-500 ng/μL.

RNA samples were submitted to the Microbial Genome Sequencing Center (MiGS) in Pittsburgh, PA. MiGS performed Illumina Stranded RNA library preparation paired with RiboZero Plus (per manufacturer’s specifications) and sequenced on a NextSeq 500 using a 75cyc High Output flowcell. Fastq files were trimmed and filtered using a combination of Trimmomatic (70) and FastQC (https://www.bioinformatics.babraham.ac.uk/projects/fastqc/). Alignment, normalization and differential expression was analyzed with SeqMan NGen® Version 17.2. DNASTAR, Madison, WI. Annotation was imported from NCBI with additional information obtained from progenomes (71). All subsequent analysis was performed in Excel and GraphPad Prism 9.

### Bioinformatics

Gene sequences and operon organization were obtained from the BioCyc data collection (35, 72). SignalP 5.0 (73) was used to search for Type I and Type II signal peptides. BUSCA (74) was used to predict protein cellular location. When all *C. difficile* proteins were run through the prediction programs, SignalP predicted ∼7% to be exported. BUSCA predicted ∼4% to exported and ∼25% to be membrane associated. Cell wall association in Tables 2 and 3 was reported as follows: (i) signal peptidase substrate if predicted by SignalP; (ii) for all remaining genes: membrane protein if predicted by BUSCA. Putative WalR binding sites were identified with Virtual Footprint (75) by querying with (1) the *B. subtilis* consensus motif, TGTWAH-N5-TGTWAH (14), (2) the same motif, but allowing one mismatch, or (3) TGTNDH-N5-BKBWRN (8). Searches were limited to the intergenic region of the genome and generated 29, 522, and 684 hits respectively.

## Data availability

RNA-seq data were submitted to the NCBI GEO repository and assigned accession number GSE200346.

## Acknowledgements

This work was supported by National Institutes of Health grants R01AI087834 (CDE) and R01AI155492 (CDE and DSW) from the National Institutes for Allergy and Infectious Disease. Cell fluorescence was quantitated at the Flow Cytometry Facility, which is a Carver College of Medicine / Holden Comprehensive Cancer Center core research facility at the University of Iowa. The facility is funded through user fees and the generous financial support of the Carver College of Medicine, Holden Comprehensive Cancer Center, and Iowa City Veteran’s Administration Medical Center. RNA integrity was characterized by the Genomics Division of the Iowa Institute of Human Genetics which is supported, in part, by the University of Iowa Carver College of Medicine. We thank Nigel Minton for pMTL-YN1C, Brianne Zbylicki for plasmid pBZ101, and members of the Ellermeier and Weiss laboratories for helpful discussions.

