## Supplemental tables S2-S4, Supplemental figures S1-S6 for "The WalRK two-component system is essential for proper cell envelope biogenesis in *Clostridioides difficile*"

Table S2 Plasmids used in this study. A. Construction details.

| Plasmid | Relevant features | Parent vector | Restriction enzymes to digest parent vector | PCR primers | PCR template | Assembly | Comments | Reference |
|---|---|---|---|---|---|---|---|---|
| pAP114 | $P_{xyl}$ ::*mCherryOpt catP* | | | | | | | (26) |
| pBZ101 | $P_{xyl}$ empty vector | pAP114 | BamHI, SacI | | | Ligation | Blunt end with Klenow, ligate with T4 DNA ligase | |
| pCE636 | $P_{dltD}$ ::*mCherryOpt catP* | pDSW1728 | KpnI, SacI | 4141+4142 | R20291 | ITA | 191 nucleotides upstream of dltD (cdr_2745) cloned before mCherry reporter | This study |
| pCE655 | $P_{xyl}$ ::*dCas9-opt* $P_{gdh}$ ::*sgRNA-pgdA-1 catP* | | | | | | | (37) |
| pCE691 | $P_{xyl}$ ::*walR catP* | pAP114 | SacI, BamHI | 4546+4547 | R20291 | ITA | *walR* from R20291 cloned under xylose control | This study |
| pCE738 | $P_{xyl}$ ::*dCas9-opt* $P_{gdh}$ ::*sgRNA-dltD-1 catP* | | | | | | | (37) |
| pCE741 | $P_{xyl}$ ::*cdr2656* | pAP114 | BamHI, SacI | 4722+4723 | R20291 | ITA | cdr_2656 (ortholog of cd630_27680) under xylose control | This study |
| pCE744 | $P_{xyl}$ ::*dCas9-opt* $P_{gdh}$ ::*sgRNA-cd630_27680-1 catP* | pIA33 | MscI, NotI | 4719+4084 | pIA33 | ITA | Replace sgRNA-rfp with sgRNA-cd630_27680-1 | This study |
| pCE745 | $P_{xyl}$ ::*dCas9-opt* $P_{gdh}$ ::*sgRNA-cd630_27680-2 catP* | pIA33 | MscI, NotI | 4720+4084 | pIA33 | ITA | Replace sgRNA-rfp with sgRNA-cd630_27680-2 | This study |
| pCE789 | $P_{xyl}$ ::*dCas9-opt* $P_{gdh}$ ::*cwpV-1 catP* | pIA33 | MscI, NotI | 4939+4084 | pIA33 | ITA | Replace sgRNA-rfp with sgRNA-cd630_cwpV | This study |
| pCE791 | $P_{xyl}$ ::*cwpV* | pAP114 | BamHI, SacI | 4937+4938 | R20291 | ITA | *cwpV* from R20291 under xylose control | This study |
| pDSW1728 | $P_{tet}$ ::*mCherryOpt catP* | | | | | | | (55) |
| pDSW2037 | $P_{xyl}$ in integration vector pMTL-YN1C *catP* | pMTL-YN1c | SacI, SalI | P2405+P2406 | R20291 | Ligation | Vector suitable for inserting genes under $P_{xyl}$ -control at *pyrE* in 630Δerm | This study |
| pDSW2053 | $P_{xyl}$ ::*dCas9opt-P* $_{gdh}$ ::*sgRNA-neg* in pMTL-YN1C *catP* | pDSW2037 | SalI+NheI | P2408+P2411 | pIA34 | Ligation | Vector suitable for integration at *pyrE* in 630Δerm, cargo: $P_{xyl}$ ::*dCas9opt-* $P_{gdh}$ ::*sgRNA-neg* | This study |
| pDSW2055 | $P_{xyl}$ ::*dCas9opt-P* $_{gdh}$ ::*sgRNA-cd630_17810-1* in pMTL-YN1C *catP* | pDSW2053 | SalI+NheI | P2408+P2411 | pIA50 | Ligation | Vector suitable for integration at *pyrE* in 630Δerm, cargo: $P_{xyl}$ ::*dCas9opt-* $P_{gdh}$ ::*sgRNA-cd630_17810-1* | This study |
| pDSW2057 | $P_{xyl}$ ::*dCas9opt-P* $_{gdh}$ ::*sgRNA-cd630_17810-2* in integration vector pMTL-YN1C *catP* | pDSW2053 | SalI+NheI | P2408+P2411 | pIA51 | Ligation | Vector suitable for integration at *pyrE* in 630Δerm, cargo: $P_{xyl}$ ::*dCas9opt-* $P_{gdh}$ ::*sgRNA-cd630_17810-2* | This study |
| pIA33 | $P_{xyl}$ ::*dCas9-opt* $P_{gdh}$ ::*sgRNA-rfp catP* | | | | | | | (26) |
| pIA34 | $P_{xyl}$ ::*dCas9-opt* $P_{gdh}$ ::*sgRNA-neg catP* | | | | | | | (26) |
| pIA50 | $P_{xyl}$ ::*dCas9-opt* $P_{gdh}$ ::*sgRNA-cd630_17810-1 catP* | pIA33 | MscI, NotI | 4454+4084 | pIA33 | ITA | Replace sgRNA-rfp with sgRNA-cd630_17810-1; will also target cdr_1676 | This study |
| pIA51 | $P_{xyl}$ ::*dCas9-opt* $P_{gdh}$ ::*sgRNA-cd630_17810-2 catP* | pIA33 | MscI, NotI | 4455+4084 | pIA33 | ITA | Replace sgRNA-rfp with sgRNA-cd630_17810-2; will also target cdr_1676 | This study |
| pIA75 | $P_{xyl}$ ::*walR* in integration vector pMTL-YN1C *catP* | pDSW2037 | HindIII, SalI | 4658+4659 | pCE691 | ITA | Vector suitable for integration at *pyrE* in 630Δerm, cargo: $P_{xyl}$ ::*walR* | This study |
| pIA76 | $P_{xyl}$ :*walR-D66E* in integration vector pMTL-YN1C *catP* | pDSW2037 | HindIII, SalI | 4658+4660, 4661+4659 | pCE691 | ITA | Vector suitable for integration at *pyrE* in 630Δerm, cargo: $P_{xyl}$ ::*walR* -D66E | This study |
| pIA79 | $P_{xyl}$ ::*dCas9-opt* $P_{gdh}$ ::*cd630_25040-1 catP* | pIA33 | MscI, NotI | 5110+4084 | pIA33 | ITA | Replace sgRNA-rfp with sgRNA-cd630_25040-1 | This study |
| pIA80 | $P_{xyl}$ ::*dCas9-opt* $P_{gdh}$ ::*cd630_25040-2 catP* | pIA33 | MscI, NotI | 5111+4084 | pIA33 | ITA | Replace sgRNA-rfp with sgRNA-cd630_25040-2 | This study |
| pIA81 | $P_{xyl}$ ::*dCas9-opt* $P_{gdh}$ ::*cd630_36010-1 catP* | pIA33 | MscI, NotI | 5112+4084 | pIA33 | ITA | Replace sgRNA-rfp with sgRNA-cd630_36010-1 | This study |

Table S2 Plasmids used in this study. A. Construction details.

| Plasmid | Relevant features | Parent vector | Restriction enzymes to digest parent vector | PCR primers | PCR template | Assembly | Comments | Reference |
|---|---|---|---|---|---|---|---|---|
| pIA93 | $P_{cdr\_0665}$ *::mCherryOpt catP* | pDSW1728 | KpnI, SacI | 5394+5395 | R20291 | ITA | 300 nucleotides upstream of cdr_0665 (ortholog of CD630_07380) plus first two codons cloned before mCherry reporter | This study |
| pIA95 | $P_{cdr\_0796}$ *::mCherryOpt catP* | pDSW1728 | KpnI, SacI | 5375+5376 | R20291 | ITA | 300 nucleotides upstream of cdr_0796 (ortholog of CD630_08670) plus first two codons cloned before mCherry reporter | This study |
| pIA97 | $P_{cdr\_2753}$ *::mCherryOpt catP* | pDSW1728 | KpnI, SacI | 5396+5397 | R20291 | ITA | 300 nucleotides upstream of cdr_2753 (ortholog of CD630_28620) plus first two codons cloned before mCherry reporter | This study |
| pIA98 | $P_{cdr\_0455}$ *::mCherryOpt catP* | pDSW1728 | KpnI, SacI | 5398+5399 | R20291 | ITA | 300 nucleotides upstream of cdr_0455 (ortholog of CD630_53000) plus first two codons cloned before mCherry reporter | This study |
| pIA100 | $P_{cd630\_0739}$ *::mCherryOpt catP* | pDSW1728 | KpnI, SacI | 5420+5421 | 630Δerm | ITA | 300 nucleotides upstream of CD630_07390 plus first two codons cloned before mCherry reporter | This study |
| pIA101 | $P_{cd630\_0739}$ *::mCherryOpt* in pMTL-YN1C *catP* | pMTL-YN1C | NotI, HindIII | 5485+5486 | pIA100 | ITA | Vector suitable for integration at *pyrE* in 630Δerm, cargo: $P_{cd630\_07390}$ *::mCherryOpt* | This study |
| pIA102 | $P_{xyl}$*::dCas9-opt* $P_{gdh}$*::cd630_05490 catP* | pIA33 | MscI, NotI | 5641+4084 | pIA33 | ITA | Replace sgRNA-rfp with sgRNA-cd630_05490 | This study |
| pIA103 | $P_{xyl}$*::dCas9-opt* $P_{gdh}$*::cd630_27940 catP* | pIA33 | MscI, NotI | 5642+4084 | pIA33 | ITA | Replace sgRNA-rfp with sgRNA-cd630_27940 (cwp12) | This study |
| pIA104 | $P_{xyl}$*::dCas9-opt* $P_{gdh}$*::cd630_27860 catP* | pIA33 | MscI, NotI | 5643+4084 | pIA33 | ITA | Replace sgRNA-rfp with sgRNA-cd630_27860 (cwp5) | This study |
| pIA105 | $P_{xyl}$*::dCas9-opt* $P_{gdh}$*::cd630_27910 catP* | pIA33 | MscI, NotI | 5644+4084 | pIA33 | ITA | Replace sgRNA-rfp with sgRNA-cd630_27910 (cwp2) | This study |
| pIA106 | $P_{xyl}$*::dCas9-opt* $P_{gdh}$*::cd630_10360 catP* | pIA33 | MscI, NotI | 5645+4084 | pIA33 | ITA | Replace sgRNA-rfp with sgRNA-cd630_10360 (cwp17) | This study |
| pIA107 | $P_{pgdA}$ *::mCherryOpt catP* | pDSW1728 | KpnI, SacI | 5664+5665 | 630Δerm | ITA | 300 nucleotides upstream of pgdA (CD630_15220) plus first two codons cloned before mCherry reporter | This study |
| pIA108 | $P_{xyl}$*::dCas9-opt* $P_{gdh}$*::cd630_07380-1 catP* | pIA33 | MscI, NotI | 5705+4084 | pIA33 | ITA | Replace sgRNA-rfp with sgRNA-cd630_380-1 | This study |
| pIA109 | $P_{xyl}$*::dCas9-opt* $P_{gdh}$*::cd630_07380-2 catP* | pIA33 | MscI, NotI | 5707+4084 | pIA33 | ITA | Replace sgRNA-rfp with sgRNA-cd630_380-2 | This study |
| pIA110 | $P_{xyl}$*::dCas9-opt* $P_{gdh}$*::cd630_07390-1 catP* | pIA33 | MscI, NotI | 5706+4084 | pIA33 | ITA | Replace sgRNA-rfp with sgRNA-cd630_390-1 | This study |
| pIA111 | $P_{xyl}$*::dCas9-opt* $P_{gdh}$*::cd630_07390-2 catP* | pIA33 | MscI, NotI | 5708+4084 | pIA33 | ITA | Replace sgRNA-rfp with sgRNA-cd630_390-2 | This study |
| pIA112 | $P_{xyl}$ *::cd630_0738* | pAP114 | BamHI, SacI | 5715+5726 | 630Δerm | ITA | cd630_0738 from 630Δerm under xylose control | This study |
| pIA113 | $P_{xyl}$ *::cd630_0739* | pAP114 | BamHI, SacI | 5717+5718 | 630Δerm | ITA | cd630_0739 from 630Δerm under xylose control | This study |
| pIA114 | $P_{xyl}$ *::cd630_0738-cd630_0739* | pAP114 | BamHI, SacI | 5715+5719, 5720+5718 | 630Δerm | ITA | cd630_0738 and cd630_0739 from 630Δerm under xylose control; separate Shine Dalgano sequence | This study |
| pIA115 | $P_{xyl}$ *::cd630_0739-cd630_0738* | pAP114 | BamHI, SacI | 5717+5721, 5722+5725 | 630Δerm | ITA | cd630_0739 and cd630_0738 from 630Δerm under xylose control; separate Shine Dalgano sequence | This study |
| pMTL-YN1C | *E. coli-C. difficile* shuttle vector for inserting genes into *C. difficile* chromosome while restoring *pyrE*; *colE1 RP4oriT-TraJ CB102ori-repH' catP* | | | | | | | (29) |

### Table S2 B. Oligonucleotides

| Oligo | Sequence | Relevant features |
|---|---|---|
| CDEP4084 | AACTTATAGGATCCGCGGCCGCTAGTCAGACATCATGCTGATCTAGA | Underlined: BamHI site |
| CDEP4141 | GGCTTCTTATTTTTATGGTACAAATTATAAGTATGAAAAAGTGCTAAAAAG | |
| CDEP4142 | TCTCCTTTACTGCAGGAGCTATTTTCTCTCCTCTAAAAATATTCAAAT | |
| CDEP4454 | AATTAAACTGTAAATGGCCA **AGTAACAAATAACATCATCA** GTTTTAGAGCTAGAAATAGC | Underlined: MscI; Bold: sgRNA-cdr_1676-1 |
| CDEP4455 | AATTAAACTGTAAATGGCCA **GGCAATATCATAGATGAATC** GTTTTAGAGCTAGAAATAGC | Underlined: MscI; Bold: sgRNA-cdr_1676-2 |
| CDEP4546 | CGATAGTTATGAAGTGAGCTTAAGGAGGATTAATATATGGTTAATATTATACTCAATTGGA | |
| CDEP4547 | GTTTTATTAAAACTTATAGGATCCAGACCTTCAAAATGTAACACT | Underlined: BamHI site |
| CDEP4658 | AGGGTAAAGAGGAGAGTCGACTAAGGAGGATTAATATATGGTTAATATTATACT | Underlined: SalI site |
| CDEP4659 | ACGACGGCCAGTGCCAAGCTTTATTCTTCCTTTCTAAAACAGTATCC | Underlined: HindIII |
| CDEP4660 | ATATCTGGAAGCATCACTTCTAAAAGAGCTATTGTAATATCATTTTCTG | |
| CDEP4661 | TTACAATAGCTCTTTTAGAAGTGATGCTTCCAGATATAAGTG | |
| CDEP4719 | AATTAAACTGTAAATGGCCA **GCACTCATAGGTGAATTAGT** GTTTTAGAGCTAGAAATAGC | Underlined: MscI; Bold: sgRNA-cd630_27680-1 |
| CDEP4720 | AATTAAACTGTAAATGGCCA **TTACATTTAAAAAATCACAC** GTTTTAGAGCTAGAAATAGC | Underlined: MscI; Bold: sgRNA-cd630_27680-2 |
| CDEP4722 | CGATAGTTATGAAGTGAGCTCGAAAGGATTAGGTATTTGAGAT | Underlined: SacI site |
| CDEP4723 | TTATTAAAACTTATAGGATCCCTATGTAAATCAATATCTATTAttaTTGTAAAACACGT | Underlined: BamHI site |
| CDEP4937 | CGATAGTTATGAAGTGAGCTCGGGAACAAAGTTAAAAAATTAAGT | |
| CDEP4938 | AAGTTTTATTAAAACTTATAGGATCGCTTTTTATTTCttaTTTCATAAATCCTAGT | |
| CDEP4939 | AATTAAACTGTAAATGGCCA **GCATTTTTTGCTTTTGCAAA** GTTTTAGAGCTAGAAATAGC | Underlined: MscI; Bold: sgRNA-cwpV |
| CDEP5110 | AATTAAACTGTAAATGGCCA **TCCTGAGGAGAATAATCTTT** GTTTTAGAGCTAGAAATAGC | Underlined: MscI; Bold: sgRNA-25040-1 |
| CDEP5111 | AATTAAACTGTAAATGGCCA **CTCGCTTCACTGCTTTCCTG** GTTTTAGAGCTAGAAATAGC | Underlined: MscI; Bold: sgRNA-25040-2 |
| CDEP5112 | AATTAAACTGTAAATGGCCA **TATAAATACTACTAAAATAA** GTTTTAGAGCTAGAAATAGC | Underlined: MscI; Bold: sgRNA-36010-1 |
| CDEP5375 | CAGGCTTCTTATTTTTATGGTACCGTATTGGTCATGTTGATGGATTTG | Underlined: KpnI site |
| CDEP5376 | TCTCCTTTACTGCAGGAGCTCTTATTTCATATATACTACCTCTCTTTTCTTTAG | Underlined: SacI site |
| CDEP5394 | CAGGCTTCTTATTTTTATGGTACCCAATCCTTCTTATTTACTTAAAATCCTG | Underlined: KpnI site |
| CDEP5395 | TCTCCTTTACTGCAGGAGCTCTTATTTCATATTTAGTTCTCCTTTTTAAATTTATTG | Underlined: SacI site |
| CDEP5396 | CAGGCTTCTTATTTTTATGGTACCGAACTTTATATCATATGTAAAAATTAAGGGAG | Underlined: KpnI site |
| CDEP5397 | TCTCCTTTACTGCAGGAGCTCTTACTTCATAACAACATCCTCCAAG | Underlined: SacI site |
| CDEP5398 | caGGCTTCTTATTTTTATGGTACCTACTTTACTAAGCTACTTCCACTC | Underlined: KpnI site |
| CDEP5399 | TCTCCTTTACTGCAGGAGCTCTTATTCcatGATTTCCTCCCCAAAAG | Underlined: SacI site |
| CDEP5420 | CAGGCTTCTTATTTTTATGGTACCGACTTTTAAATTTCTTATCATAGGATTTTAAG | Underlined: KpnI site |
| CDEP5421 | TCTCCTTTACTGCAGGAGCTCTTATTTCATATCTTACCTCCTAAATTAAATG | Underlined: SacI site |
| CDEP5485 | AGGAATTAGGGATGTAATAAGCGGATCCTTATTTATATAATTCATCCATAC | |
| CDEP5486 | ACGACGGCCAGTGCCAAGCTGGTACCGACTTTTAAATTTCTTATC | |
| CDEP5641 | AATTAAACTGTAAATGGCCA**TACCATCATTGCACTTATAA** GTTTTAGAGCTAGAAATAGC | Underlined: MscI; Bold: sgRNA-05490 |
| CDEP5642 | AATTAAACTGTAAATGGCCA**GTTATAAAAACCTTATCTAT** GTTTTAGAGCTAGAAATAGC | Underlined: MscI; Bold: sgRNA-27940 |
| CDEP5643 | AATTAAACTGTAAATGGCCA **GCTCCTTTAGCCTTTGCAAA** GTTTTAGAGCTAGAAATAGC | Underlined: MscI; Bold: sgRNA-27860 |
| CDEP5644 | AATTAAACTGTAAATGGCCA **GTAGTTTCTGCAGCAAAAAC** GTTTTAGAGCTAGAAATAGC | Underlined: MscI; Bold: sgRNA-27910 |

| Oligo | Sequence | Relevant features |
|---|---|---|
| CDEP5645 | AATTAAACTGTAAATGGCCA **TCAGTAGCAAATGCCATTGT** GTTTTAGAGCTAGAAATAGC | Underlined: MscI; Bold: sgRNA-10360 |
| CDEP5664 | CAGGCTTCTTATTTTTATGGTACCCTAAATTTACATACATTTTGAGCTC | Underlined: KpnI site |
| CDEP5665 | TCTCCTTTACTGCAGGAGCTCTTATACCATAATTACAGCATTCCCC | Underlined: SacI site |
| CDEP5705 | AATTAAACTGTAAATGGCCA **AATTTACATCCATATAATCA** GTTTTAGAGCTAGAAATAGC | Underlined: MscI; Bold: sgRNA-07380-1 |
| CDEP5706 | AATTAAACTGTAAATGGCCA **ATTGCATTTTTTTCTGCTTT** GTTTTAGAGCTAGAAATAGC | Underlined: MscI; Bold: sgRNA-07390-1 |
| CDEP5707 | AATTAAACTGTAAATGGCCA **AATTATCTTGCAATTTTTCT** GTTTTAGAGCTAGAAATAGC | Underlined: MscI; Bold: sgRNA-07380-2 |
| CDEP5708 | AATTAAACTGTAAATGGCCA **AAAGTAAAAAAATTTAAAAG** GTTTTAGAGCTAGAAATAGC | Underlined: MscI; Bold: sgRNA-07390-2 |
| CDEP5715 | CGATAGTTATGAAGTGAGCTCAAGGAGGAACTAAATatgAAAAAAAGATTAATAATAATG | Underlined: SacI site |
| CDEP5717 | CGATAGTTATGAAGTGAGCTCAAGGAGGGGTAAGATatgAAAGTAAAAAAATTTAAAAG | Underlined: SacI site |
| CDEP5718 | TCTATTTAAAGTTTTATTAAAACTTATAGGATCCttaTTTTTCAAGCTCATCAAAGTAAT | Underlined: BamHI site |
| CDEP5719 | TTCATATCTTACCCCTCCTTTTTAAAGTTATTATTTCATAAGTTTATCG | |
| CDEP5720 | TAATAACTTTAAAAAGGAGGGGTAAGATATGAAAGTAAAAAAATTTAAAAG | |
| CDEP5721 | TTCATATTTAGTTCCTCCTTTCTTTAAGCATTATTTTTCAAGCTC | |
| CDEP5722 | TAATGCTTAAAGAAAGGAGGAACTAAATATGAAAAAAAGATTAATAATAATGATATTG | |
| CDEP5725 | TTATTAAAACTTATAGGATCCttaTTTCATAAGTTTATCGTTTTCCAT | Underlined: BamHI site |
| CDEP5726 | TTATTAAAACTTATAGGATCCttaTTTCATAAGTTTATCGTTTTCC | Underlined: BamHI site |
| P2405 | GCCGCTGTATCCATATGACC | |
| P2406 | AATTAGGATAGGAAAACGATAG | |
| P2408 | GAGGAGAGTCGACGCATGG | Underlined: SalI site |
| P2411 | GTGGCTAGCCTGGCATCTTTTTATTTAGGGATTTCTCAC | Underlined: NheI site |

**Table S3. Wal regulon genes tested for effects on induction of the regulon**

| Gene locus | Gene name | Description | Wal-ON or Wal-OFF | Rationale | Overexpression | CRISPRi |
|---|---|---|---|---|---|---|
| *cd630_05140* | *cwpV* | Cell wall binding protein | OFF | Most highly induced, S-layer associated | pCE791 | pCE789 |
| *cd630_05490* | | Hypothetical protein | ON | Highly induced | | pIA102 |
| *cd630_07380* | | Hypothetical protein | ON | Highly induced; lysozyme regulon[1] | pIA112, pIA114, pIA115 | pIA110, pIA111 |
| *cd630_07390* | | Hypothetical protein | ON | Most highly induced, lysozyme regulon[1] | pIA113, pIA114, pIA115 | pIA108, pIA109 |
| *cd630_10360* | *cwp17* | Cell wall binding protein | ON | Moderately induced, S-layer associated (amidase) | | pIA106 |
| *cd630_15220* | *pgdA* | PG deacetylase | ON | Highly induced, PG related | | pCE655 |
| *cd630_25040* | | D-alanyl-D-alanine carboxypeptidase | OFF | Weakly repressed, PG associated[2] | | pIA79, pIA80 |
| *cd630_27680* | | Cell-wall hydrolase | ON | Moderately repressed, hydrolase | pCE741 | pCE744 |
| *cd630_27860* | *cwp5* | Cell wall binding protein | ON | Moderately induced, S-layer associated | | pIA104 |
| *cd630_27910* | *cwp2* | Cell wall binding protein | ON | Moderately induced, S-layer associated, putat. WalR binding site | | pIA105 |
| *cd630_27940* | *cwp12* | Cell wall binding protein | ON | Moderately induced, S-layer associated, putat. WalR binding site | | pIA103 |
| *cd630_28540* | *dltD* | D-alanine transferase | OFF | Moderately repressed, cell wall associated, putative WalR binding site | | pCE738 |
| *cd630_36010* | | D-alanyl-D-alanine carboxypeptidase | ON | Moderately induced, PG related | | pIA81 |

[1]: (43)
[2]: See Table S1

**Table S4. Putative WalR binding sites**

| Gene | Name | Description | Orientation[1] | Search[2] | Wal-ON or Wal-OFF |
|---|---|---|---|---|---|
| *cd630_08670* | | TGTTTA-GTAAT-GTGAAT | Forward | C | ON, up |
| *cd630_07400* | | TGTAAA-GCCTC-CTTAAA | Reverse | C | ON, up |
| *cd630_27940* | *cwp12* | TGTGAT-TTTCA-TTTTAT | Reverse | C | ON, up |
| *cd630_03910* | *asnB* | TGTTAT-AACAC-TGATAA | Forward | B | ON, up |
| *cd630_14960* | | TGTTAT-AGAAT-CGGAAG | Reverse | C | ON, up |
| *cd630_27910* | *cwp2* | AGTAAC-AAAAT-TGTAAT | Reverse | B | ON, up |
| *cd630_28620* | | TGTTTT-GCCTT-TTTAGT | Reverse | C | ON, up |
| *cd630_20580* | | TGTAAA-TAAAC-TGTAAA | Reverse | A | ON, up |
| *cd630_07410* | *glpK1* | TGTAAA-GCCTC-CTTAAA | Forward | C | ON, up |
| *cd630_26810* | | TGTCAA-ACACA-CTGAAA | Forward | C | ON, up |
| *cd630_22280* | | TGTTTT-TTGTT-TTTTAT | Reverse | C | ON, up |
| *cd630_18070* | | TGTAAT-AAAAT-TGAAAC | Reverse | B | ON, up |
| *cd630_30360* | | TGTAAA-ACAAT-TTGTAA | Reverse | B,C | ON, down |
| *cd630_10540* | *bcd2* | TGTCAT-AATTG-GTTAGT<br>TGTTGC-CGAGA-TTTTAA | Forward<br>Forward | C<br>C | ON, down |
| *cd630_27680* | | TGTAAT-CTTTT-TGTAAT<br>TGTAAT-CTTTT-TGTAAT | Reverse<br>Reverse | A<br>A | ON, down |
| *cd630_22600* | | TGGTAA-AATTT-TGTTAA | Forward | B | ON, down |
| *cd630_17290* | | TGTAAA-AAATA-TGTAAA | Reverse | A | Off, up |
| *cd630_23920* | | TGTTTA-AACTA-TTGAAA | Reverse | C | Off, up |
| *cd630_32630* | *pstC* | TGTATT-CAAAA-TTTAAT | Reverse | C | Off, up |
| *cd630_28540* | *dltD*[3] | TGTTAC-AAAAT-TGTAAA | Reverse | A | Off, down |
| *cd630_05300* | | TGTTTA-AATAT-TGTAGG | Reverse | C | Off, down |
| *cd630_28620* | | TGTTTT-GCCTT-TTTAGT | Reverse | C | Off, down |

[1]:Orientation: Putative WalR binding site is in the same orientation as gene (forward) or in the opposite direction (reverse).
[2]:Search catergories: (A) TGTWAH-N5-TGTWAH (14), (B) ) TGTWAH-N5-TGTWAH, one mismatch allowed, (C) TGTNDH-N5-BKBWRN (8).
[3]: Previously described as part of a tandem direct repeat sequence (49).

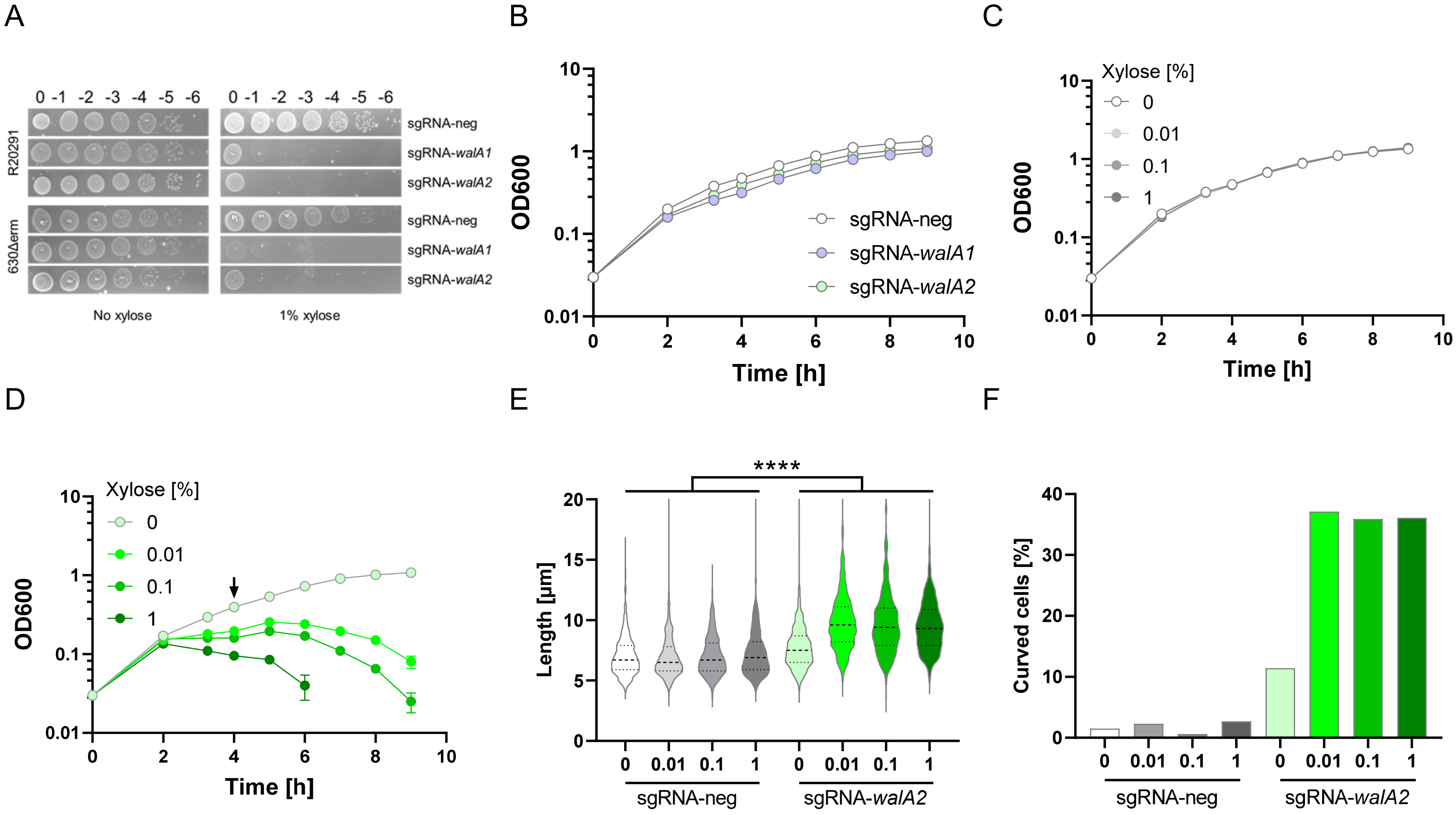


**Fig. S1. CRISPRi silencing of the *wal* operon.** (A) Plasmid-based CRISPRi silencing of the *wal* operon causes a strong viability defect. Serial dilutions of overnight cultures were spotted onto TY-Thi plates with or without 1% xylose to induce the expression of dCas9. Plates were photographed after incubation overnight. The strains shown are exconjugants of R20291 or 630Δerm harboring the following CRISPRi plasmids: pIA34 (sgRNA-neg, negative control), pIA50 (sgRNA-*walA1*) or pIA51 (sgRNA-*walA2*). (B-F). Silencing of the *wal* operon with a chromosomal copy of the CRISPRi machinery causes growth and morphology defects. (B) Growth curves of sgRNA-neg (UM554), sgRNA-*walA1* (UM555) and sgRNA-*walA2* (UM556) in TY, no xylose. (C) Growth curve of sgRNA-neg (UM554) in TY with 0, 0.01, 0.1 or 1% xylose. (D) Growth curves of sgRNA-*walA2* (UM556) in TY with 0, 0.01, 0.1 or 1% xylose. (E) Cell length and (F) percent cells with curved contours based on measurements of about 500 cells per condition. Samples were taken 4h after subculture (arrow in D). The sgRNA-*walA2* strain is longer than the sgRNA-neg control in all pairwise comparisons as determined by a T-test, $p<0.0001$. Data shown are representative of at least 2 independent experiments.

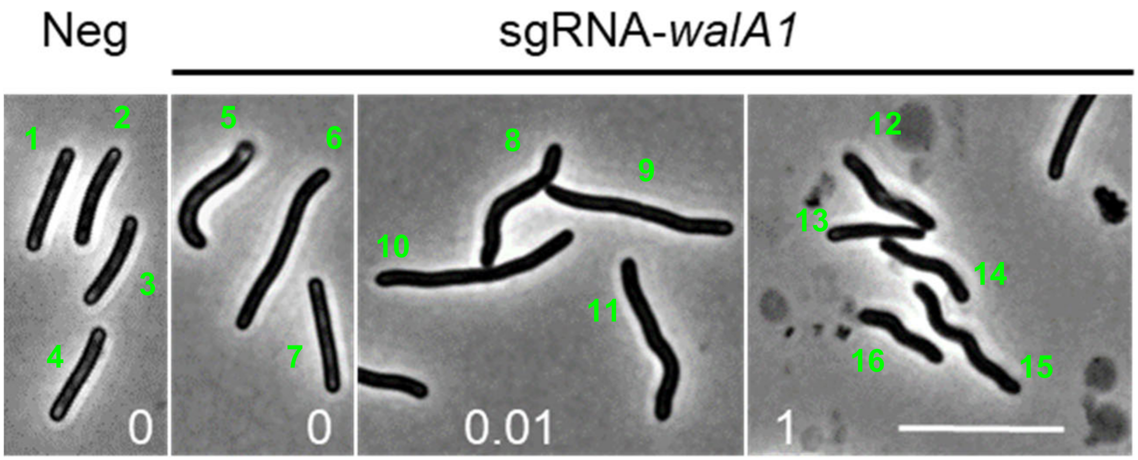


| Cell | Sinuosity |
|---|---|
| 1 | 1.001 |
| 2 | 1.008 |
| 3 | 1.010 |
| 4 | 1.003 |
| 5 | 1.183 |
| 6 | 1.011 |
| 7 | 1.001 |
| 8 | 1.066 |

| Cell | Sinuosity |
|---|---|
| 9 | 1.009 |
| 10 | 1.032 |
| 11 | 1.046 |
| 12 | 1.020 |
| 13 | 1.008 |
| 14 | 1.039 |
| 15 | 1.039 |
| 16 | 1.032 |

**Fig. S2. Sinuosity examples.** Cells from Fig 2C are numbered and their measured sinuosity reported in the table below. Orange highlighting indicates sinuosity values above 1.03, i.e. cells that were counted as "curved" in Fig. 2E. Strains shown are sgRNA-neg (UM554) and sgRNA-*walA1* (UM555). Numbers at the bottom of each image are the % xylose used for induction of *dCas9*. Bar = 10 µm.

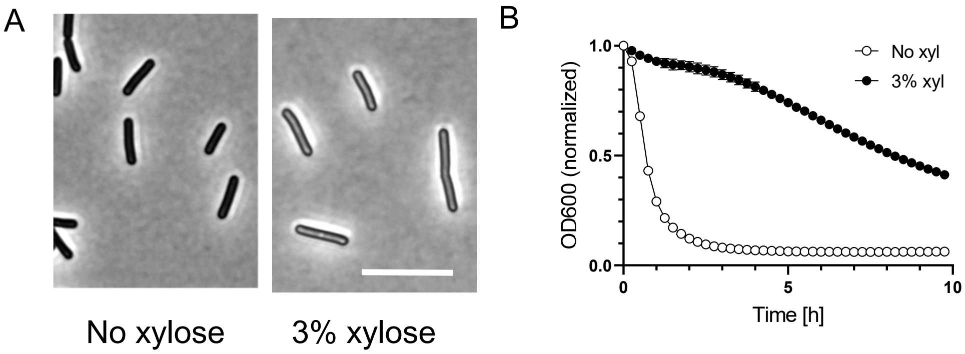


**Fig. S3. Overexpression of *walR* from a multi-copy plasmid alters morphology and slows autolysis in R20291**. (A) Phase contrast microscopy and (B) Lysis assay of R20291 harboring pCE691 ($P_{xyl}$::*walR*). Duplicate cultures were grown in TY-Thi to an $OD_{600}$ of 0.2, at which time one was induced with 3% xylose (green arrow). Both cultures were harvested after 3h. Bar = 10 µm. Data shown are representative of at least 3 experiments. Error bars in (B) depict SD of 3 technical replicates but are mostly smaller than the symbols.

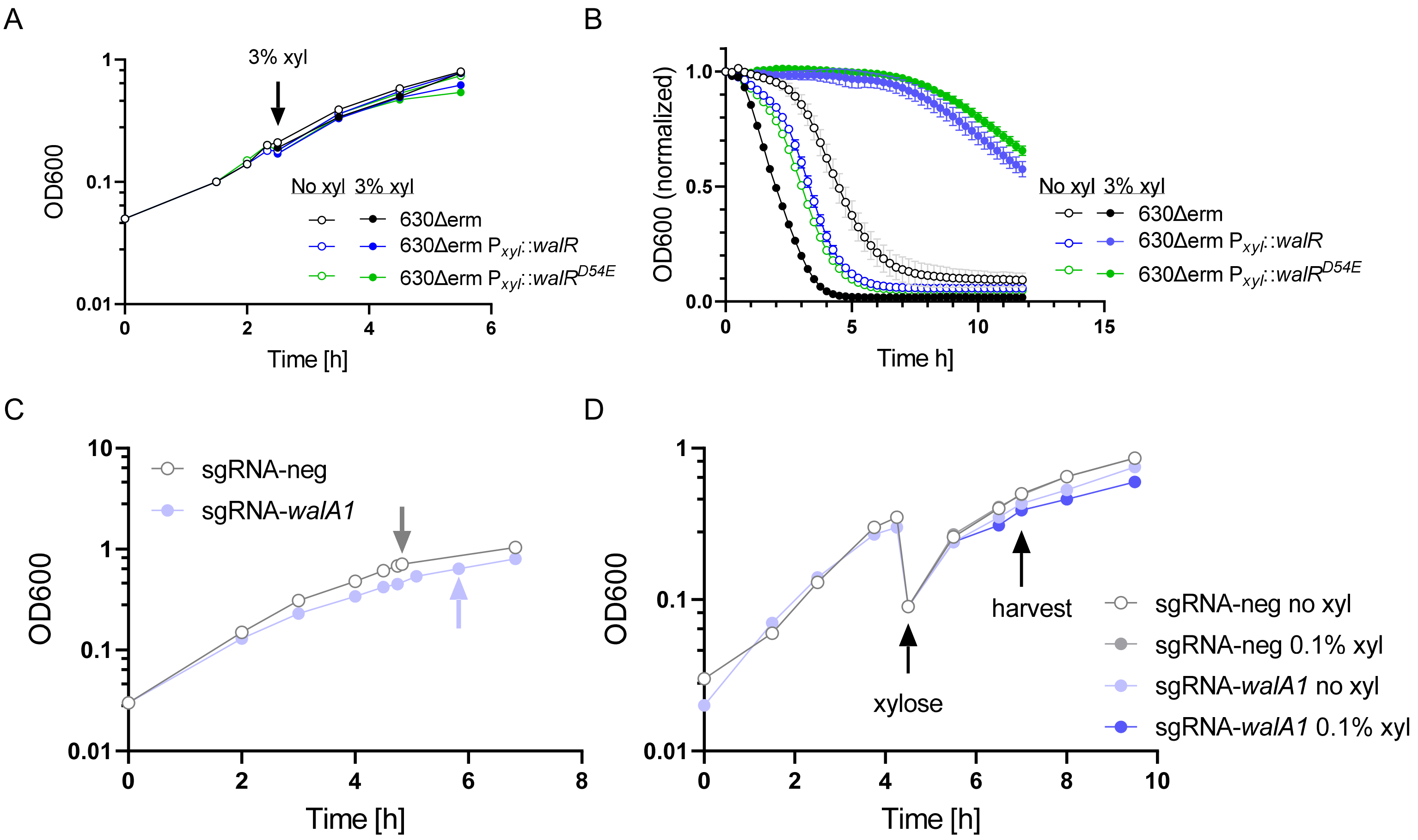


**Fig. S4. Culture growth for RNA-seq experiments**. (A, B) $P_{xyl}$::*walR* strains for RNA-seq under Wal-ON conditions. (A) Growth curves. (B) Lysis assay. Error bars in (B) depict SD of 3 technical replicates but are mostly smaller than the symbols. Strain 630Δerm (UM275) and derivatives with $P_{xyl}$::*walR* (UM626) or $P_{xyl}$::*walR-D54E* (UM628) integrated at *pyrE* were grown overnight in TY-Thi. These cultures were diluted in duplicate into the same medium to a starting $OD_{600}$ of 0.05, grown to $OD_{600}$ ~0.2, one culture set was induced with 3% xylose, and all cultures were harvested 3h later. RNA was purified from the induced cultures only. The non-induced cultures were included for growth and lysis phenotype comparison. (C, D) sgRNA-*walA1* strain for RNA-seq under Wal-OFF conditions. (C) Growth curves in TY without xylose. Overnight cultures of the CRISPRi strain UM555 (sgRNA-*walA1*) and the matched control UM554 (sgRNA-neg) were subcultured to $OD_{600}$ of 0.03 and harvested at $OD_{600}$ ~0.7 (arrows). (D) Growth curves of the same strains in TY with low xylose induction. Overnight cultures grown in TY were subcultured into TY to $OD_{600}$ of 0.03 and grown to $OD_{600}$ of 0.3. These cultures were then diluted in duplicate to $OD_{600}$ = 0.1 and one culture set was induced with 0.1% xylose. Cells were harvested after 2.5h of induction. RNA was purified from the induced cultures only. The non-induced cultures were included for growth comparison.

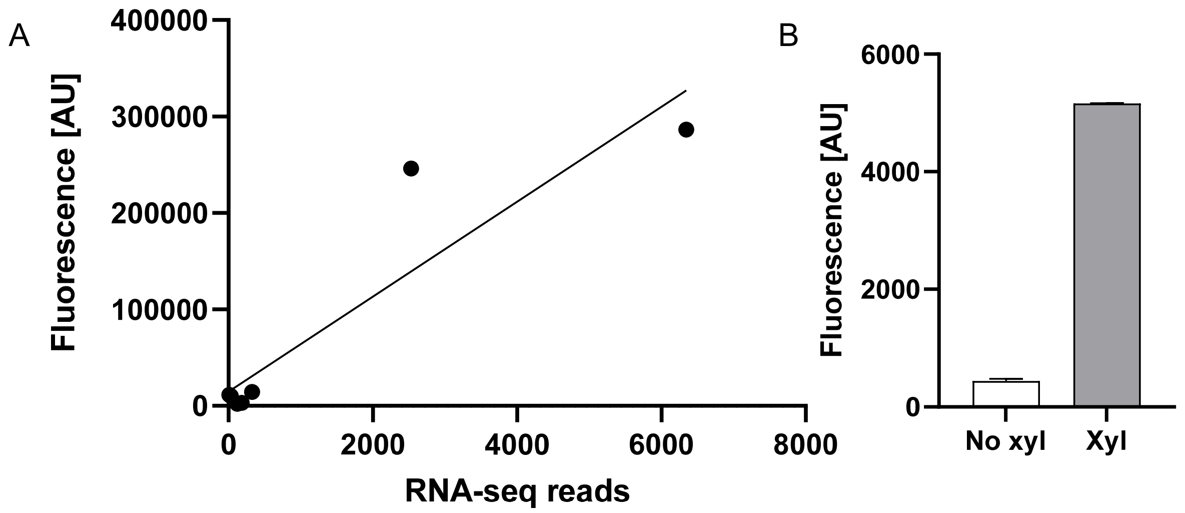


**Fig. S5. RNA-seq follow-up.** (A) Baseline expression of *rfp* reporter plasmids correlates with baseline RNA-seq reads. 630Δerm exconjugants harboring plasmids with transcriptional fusions of *rfp* to several WalR regulon promoters were grown in TY-Thi and red fluorescence was measured by flow cytometery. RNA-seq reads are from strain UM554 grown in TY without xylose. The genes shown are the same as Fig. 6: *pgdA, dltD, cd630_0738, cd630_0739, cd630_08670, cd630_ 53000, cd630_28620.* (B) Chromosomal $P_{cd630\_0739}$*::rfp* responds to overexpression of walR. Strain UM926 (630Δerm $P_{cd630\_0739}$*::rfp*) harboring plasmid pCE691 ($P_{xyl}$::walR) was subcultured to $OD_{600}$ of 0.05, grown to 0.35 and induced with 3% xylose for 3 h. Fluorescence was measured by flow cytometry.

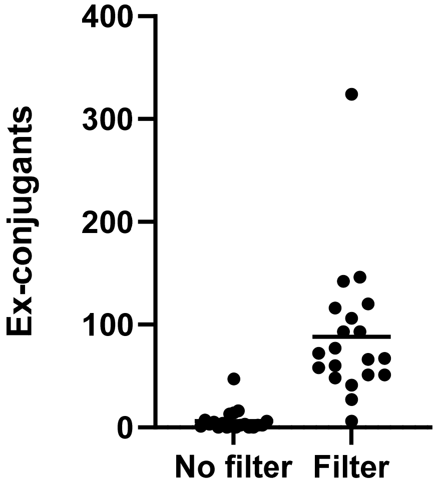


**Fig. S6. Mating on filters improves conjugation efficiency.** Twenty different CRISPRi plasmids were conjugated from *E. coli* into R20291 by mating directly on BHI plates or on cellulose filters. Conjugation success reported as number of ex-conjugants increased by more than 10-fold with the use of filters.
